# Unravelling the conundrum of nucleolar NR2F1 localization: A comparative analysis of NR2F1 antibody-based approaches *in vitro* and *in vivo*

**DOI:** 10.1101/2023.12.31.573551

**Authors:** Michele Bertacchi, Susanne Theiß, Ayat Ahmed, Michael Eibl, Agnès Loubat, Gwendoline Maharaux, Wanchana Phromkrasae, Krittalak Chakrabandhu, Aylin Camgöz, Christian Schaaf, Michèle Studer, Magdalena Laugsch

## Abstract

As a transcription factor, *NR2F1* regulates spatiotemporal gene expression during development and in adulthood. Aberrant *NR2F1* causes a rare neurodevelopmental disorder known as Bosch-Boonstra-Schaaf Optic Atrophy Syndrome. In addition, altered *NR2F1* expression is frequently observed in various cancers and is considered a prognostic marker or potential therapeutic target. In this context, NR2F1 has been shown to localize not only in the nucleus but also in the nucleoli, suggesting a novel non-canonical role in this compartment. Hence, we studied this phenomenon employing various *in vitro* and *in vivo* models in different antibody-dependent approaches. Examination of seven commonly used anti-NR2F1 antibodies in different human cancer and stem cells as well as in wild type and null mice revealed that the nucleolar localization of NR2F1 is artificial and does not play a functional role. Our subsequent comparative analysis demonstrated for the first time which anti-NR2F1 antibody best fits which approach. As our data allow for correct data interpretation, making them publicly available may have far-reaching implications for NR2F1 research in health and disease. More generally, the study also underlines the need to optimize any antibody-mediated technique.

## INTRODUCTION

Studying the spatial abundance and role of proteins under physiological and pathological conditions requires specific antibodies (Abs) to detect and quantify these proteins by widely used methods such as Western blot (WB) and flow cytometry (FC) or immunofluorescence (IF). False-positive findings, e.g., due to unspecific staining and false-negative findings, due to the inability to detect an antigen for methodological reasons such as fixation or tissue processing can have serious implications for data interpretation^1–3^. NR2F1 (COUP-TFI), a transcription factor (TF) of the steroid/thyroid hormone receptor superfamily, is no exception in this regard, as multiple Ab-based experimental approaches have been used in the past to characterize its physiological expression pattern and the pathological consequences of its perturbation. In humans, *NR2F1* haploinsufficiency caused by deletion or loss-of-function mutations results in a monogenic neurodevelopmental disorder named Bosch-Boonstra-Schaaf Optic

Atrophy Syndrome (BBSOAS)^4–6^. Characterized by intellectual disability, developmental delay, and visual impairment^3, 19, 22^, BBSOAS is often associated with additional features, such as epilepsy, autistic traits, hypotonia, oromotor dysfunction, craniofacial malformations, and hearing impairment^4–20^. *NR2F1* is now listed among the 10 top genes causally linked to rare hereditary optic neuropathies^21^. In addition, *NR2F1* has recently been linked to Hirschsprung disease^22^ and Waardenburg syndrome type IV^23^, as well as to the development of lymphoid and cardiac cells^24^, but its precise role in these disorders remains to be investigated.

The NR2F1 protein sequence is highly conserved in evolution^25^ and consists of a DNA-binding domain (DBD) to recognize its DNA binding motifs, of a hinge region (also called the D region), and of a ligand-binding domain (LBD) necessary for dimerization and co-factor binding^26–28^. Other domains pivotal for NR2F1 function are the activating function (AF) domains 1 and 2^29^, located at the N- and C-terminus, respectively. The high degree of sequence conservation, especially between human and mouse NR2F1/Nr2f1 (Nr2f1 carries the same domains, just shifted by a few amino acids), strongly suggests their evolutionary conserved functions. Studies in human cells and animal models, particularly in the mouse, have helped to characterize the expression pattern and role of human *NR2F1* and of its murine homologue *Nr2f1* in embryonic development^26, 30, 31^. NR2F1/Nr2f1 has been shown to regulate proliferation, migration, differentiation, and regional identity acquisition of neural progenitors and neurons^26, 27, 32^ (NP/N) in the developing neocortex^33, 34^ and visual system^35, 36^, and controls the balance between excitatory and inhibitory neurons^37, 38^. In addition, NR2F1 plays a crucial role in development of neural crest, a multipotent and largely embryonic cell population pivotal for e.g., craniofacial morphogenesis^39–41^. Beyond developmental stages, *Nr2f1* expression in the postnatal and adult mouse brain has been linked to neuronal activity^42–44^. NR2F1 also plays an important role in adult non-neuronal tissues, such as hematopoietic cells^45^, and its aberrant expression has been linked to cancer, serving as a prognosis marker and making it a good candidate for therapeutic development^46–52^.

Consistent with its molecular function as a TF that binds to thousands of targets including genes and their enhancers^38, 39, 41, 53^, both human NR2F1 and mouse Nr2f1 were detected by immunostaining largely within the nucleoplasm^27, 30, 31, 35, 54, 55, 56–59^. Nevertheless, some previous studies reported a cytoplasmic or nucleolar localization^45, 60–62^ raising the intriguing possibility that NR2F1 could play an additional noncanonical role in these compartments. Nucleolar aggregates observed after NR2F1 staining of tumor cells with the mouse the monoclonal Ab (clone H8132) have been used to quantify NR2F1-positive cells^60^, but the specific role of this TF in nucleoli (membraneless organelles formed by phase separation essential for ribosome biogenesis^66, 67^) remains unclear.

To explore the possible function of NR2F1 in nucleoli, we employed human neural crest cells (hNCC) derived from human induced pluripotent stem cells (hiPSC) and standard IF protocols. Using the monoclonal Ab (clone H8132), one of the most common anti-NR2F1 antibodies, we were able to confirm the nuclear and nucleolar localization of NR2F1, but the nucleolar-like pattern was also seen in cells that do not express *NR2F1* (e.g., hiPSC). Building on our experience in different methods^30, 53, 68–70^, we therefore systematically examined this Ab along with six other NR2F1 Abs using different technical set- ups (IF, FC, WB) in different cancer, wild type (*WT*), and CRISPR/Cas9 engineered stem cells, as well as in *WT* and *null* mice. We could not confirm a nucleolar-like pattern of NR2F1 in IF with any of the other Abs and our results strongly suggest that the nucleolar-like staining by a mouse monoclonal Ab is unspecific and may depend on *NR2F1* expression level, fixation methods, and tissue type.

It has long been known that establishing optimal conditions and negative controls for Ab use is critical to verify the reliability of results. This is especially true for future IF experiments with the monoclonal Ab (clone H8132) to draw correct conclusions about the role of NR2F1 in health and, if altered, its contribution to human disease.

## RESULTS

### Nucleolar-like localization of endogenous NR2F1 in hNCC observed in IF by the monoclonal Ab (clone H8132)

NR2F1 plays an essential role upon embryonic development and serves as a differentiation marker for various cell lineages, such as NP/N and hNCC^39, 57, 71–73^. Hence, to investigate the expression levels and localization of NR2F1, we used hNCC derived from healthy hiPSC employing a well-established protocol^41, 69, 74^. We performed IF with one of the commonly used NR2F1 monoclonal antibodies (clone H8132). First, we tested three different concentrations (2 µg, 1 µg and 0.4 µg/ml) of this Ab (Supplementary Figure 1A) and observed signals that were not only diffusely distributed throughout the nucleus but also in aggregate-like clusters. The data are consistent with our previous observations^69^ and have been reported in human malignant cells^61^, prostate cancer cells^62, 75^, and breast cancer cells^60^ and it is noteworthy that the same Ab was used in all these studies. While the highest Ab concentration resulted in additional staining within the cytoplasm and the lowest mostly of the aggregates, we decided to use for further experiments the intermediate concentration of 1.0 µg/ml, which showed both the diffuse nuclear signal and the nuclear aggregates (Supplementary Figure 1A). As a negative control, we used hNCC stained with either primary Ab H8132 (1°Ab) or secondary Ab (2°Ab), an anti-mouse Alexa Fluor 488 (AF488) alone or, to capture possible autofluorescence, cells stained only with DAPI to visualize the nuclei (Supplementary Figure 1B). We then tested whether the aggregates stained by the anit-NR2F1 Ab H8132 were specific to NR2F1 or also formed by some known NR2F1 co-operators. Cooperative DNA binding of different master TFs at specific loci and their synergistic role in regulating gene transcription in a spatiotemporal manner are crucial for developmental processes and non-coding elements, such as enhancers, which typically harbor multiple TF binding sites^33^. Consistent with this, chromatin binding sites occupied by NR2F1 in hNCC are frequently co-bound by the hNCC master regulator TFAP2A^39^, as shown by chromatin immunoprecipitation coupled with DNA sequencing (ChIP-seq) performed using Ab H8132, which has been widely used for this approach^33, 35^ (Supplementary Figure 1C). Hence, we tested the co- localization of both TFs by co-IF staining (Supplementary Figure 1D). Although both TFs were diffusely distributed in the nucleoplasm, only NR2F1 and not TFAP2A exhibited the aggregate-like pattern, suggesting that the aggregates may be specific for NR2F1. According to histone modifications profiles associated with active (e.g., Histone-3-lysine-27-acetylation, H3K27ac) or repressive chromatin state (e.g., Histone-3-lysine-27-trimethylation, H3K27me3), enhancers (Histone-3-lysine-4- monomethylation, H3K4me1) or promoters (Histone-3-lysine-4-trimethylation, H3K4me3), NR2F1 is associated with active enhancers in hNCC^39, 41^ (Supplementary Figure 1C). Thus, to provide further clues as to what role these NR2F1 aggregates might have, and considering that NR2F1 can activate or inactivate gene expression^25, 27, 28^, we stained NR2F1 together with H3K27ac, which marks active chromatin, and heterochromatin protein 1 (HP1), which marks an inactive chromatin state^76^. However, none of these co-stainings indicated a convincing co-localization with NR2F1 (Figure 1A). Since the NR2F1 aggregates were located in DAPI-depleted regions (Supplementary Figure 1A and D), we hypothesized that they localize to nucleoli, raising the interesting question of whether NR2F1 might be involved in ribosome biogenesis or in cellular stress response. To explore this potentially novel role of NR2F1, we performed co-staining with the nucleolar marker ZSCAN1 (zinc finger and SCAN domain containing 1)^77^ and revealed their co-localization (Figure 1B). Of note, in vertebrates, in addition to NR2F1, its homologue NR2F2 has been identified, whose nucleolar localization has been shown in breast cancer but not explored^78^. The expression patterns and functions of NR2F1 in hNCC partially overlap with those of NR2F2, which controls some of the same but also distinct genes^39, 79, 80^. Moreover, similar cooperativity as previously shown for NR2F1 and TFAP2A in hNCC was also demonstrated for NR2F1 and NR2F2^39^. However, ChIP-seq profiles lack data about TF binding to ribosomal RNA gene chromatin typically associated with nucleoli, since it is widely excluded from standard genome assemblies due to its repetitive nature^81^. As we further proved that neither TFAP2A nor NR2F2 localized to nucleoli in co-stainings with ZSCAN1 (Figure 1C and Supplementary Figure 1E), our results suggest that the nucleolar-like localization in hNCC is restricted to NR2F1.

**Figure 1:**
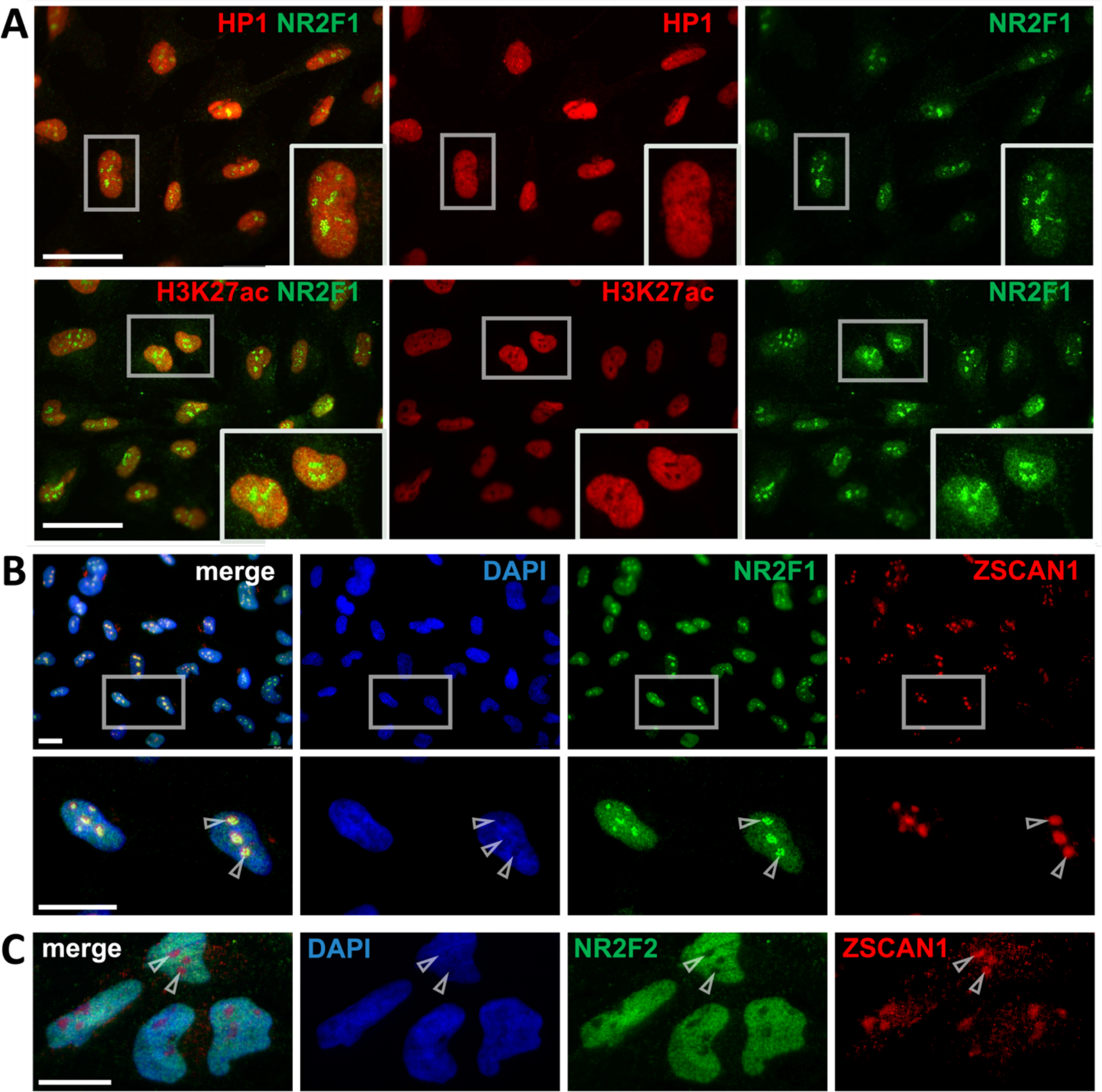
IF of endogenous NR2F1 in hiPSC-derived hNCC using Ab H8132. A) Co-IF of NR2F1 (green) either with H3K27ac (red) corresponding to active chromatin or with HP1 (red) corresponding to inactive chromatin status. B) Co-IF of NR2F1 (green) with ZSCAN1 (red) and their co-localization in nucleoli. C) Co-IF of NR2F2 (green) with ZSCAN1 (red) showed no nucleolar-like localization. Nucleoplasm (blue) was stained with DAPI. Scale bars: 20 µm.

### Evaluation of nucleolar-like localization of NR2F1 with Ab H8132 indicates an unspecific staining

The apparent co-localization of NR2F1 aggregates with ZSCAN1 in nucleoli implies that NR2F1 may be involved e.g., in ribosome biology, in addition to its canonical role as a TF. In fact, recent studies showed that a large number of nucleolar proteins are localized to multiple compartments, including the nucleoplasm and that the nucleolar proteome is much larger than previously recognized^82, 83^. However, after examination of the Human Protein Atlas (https://www.proteinatlas.org/ENSG00000175745-NR2F1/cell)^82–84^, we found no evidence of nucleolar localization of NR2F1. We also found no nucleolar NR2F1 localization when searching the Nucleolar Proteome Database (NOPdb)^85^ or in the available experimental localization data in the UniProtKB/Swiss-Prot database (The UniProt Consortium, 2019). Because in human prostate cancer samples, the nucleolar localization of NR2F1 was observed in epithelial but not in stromal cells^75^, we hypothesized that the nucleolar localization of NR2F1 might be hNCC-specific, especially since the data from hNCC was not included in the databases. To test this, we decided to use additional cell lines for further IF experiments. Since NR2F1 is highly expressed in HeLa cells (Human Protein Atlas and previous studies^86^) but not present in nucleoli (HeLa proteomics data^87^), we did not expect nucleolar localization of NR2F1 in this cell line. Given that NR2F1 is not expressed in undifferentiated pluripotent cells^88^, our undifferentiated hiPSC served as a negative control (Supplementary Figure 2.A). In addition to Ab H8132, we also used another Ab in this experiment (#6364, Cell Signaling) targeting a different region of NR2F1. As shown in Figure 2A, Ab H8132 detected the nucleolar-like aggregates appearing as in all tested cell types, including the negative control, hiPSC. In contrast, Ab #6364 stained the nucleoplasm only in HeLa and hNCC, but not in hiPSC, and foci were not observed in any of the cells. To prove whether the NR2F1-foci observed in undifferentiated hiPSC by Ab H8132 staining were still localized to nucleoli, we again performed a co-staining of NR2F1 or NR2F2 with the nucleolar marker ZSCAN1. Although, both NR2F1 (Supplementary Figure 2.A) and NR2F2^69^ are not expressed in undifferentiated hiPSC, NR2F1, but not NR2F2, showed nucleolar pattern that co-localize with ZSCAN1 (Supplementary Figure 2.B).

**Figure 2:**
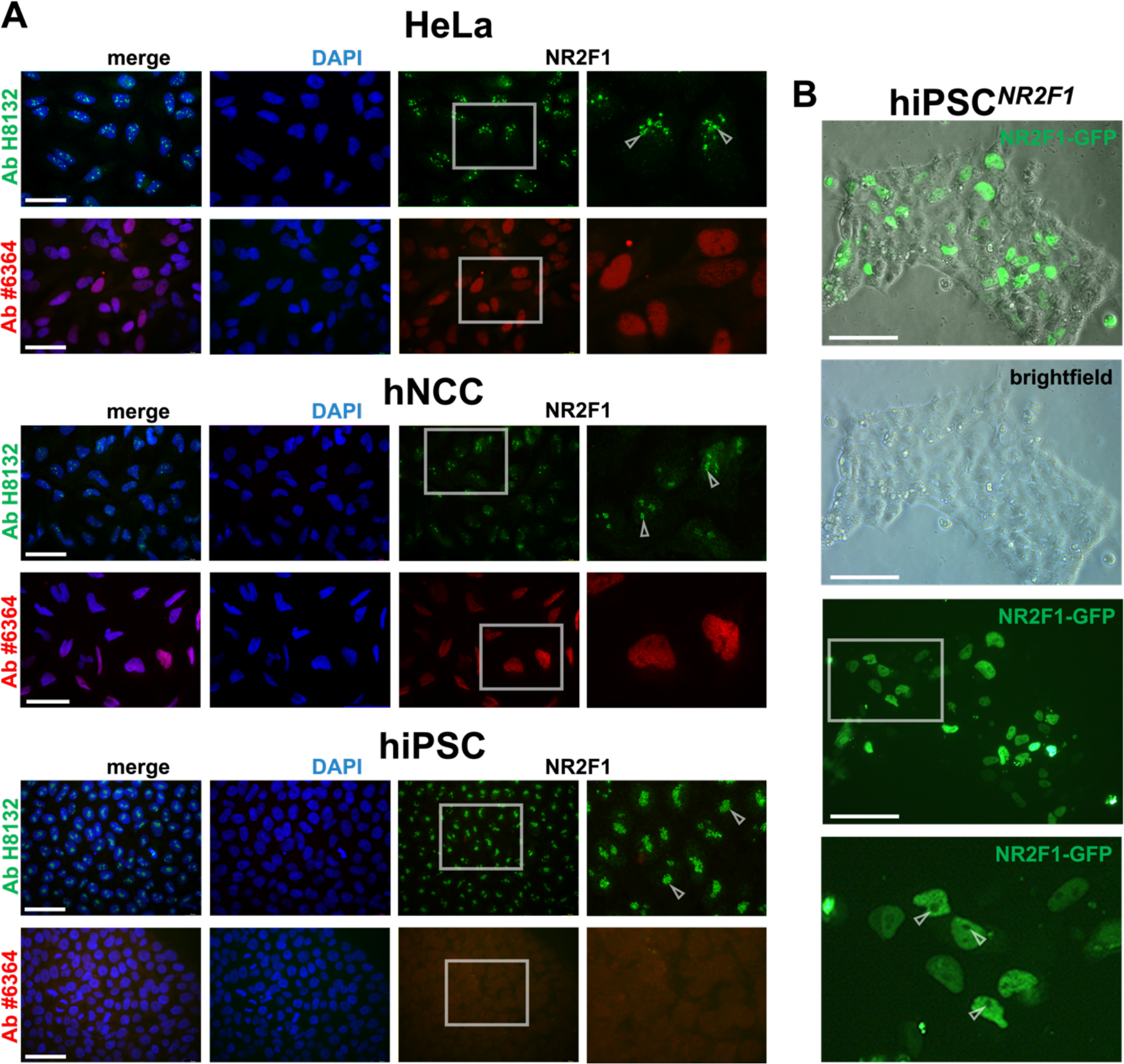
Ab-dependent NR2F1 localization tested in different cell types. A) IF of NR2F1 by Ab H8132 (green), and by Ab #6364 (red) performed in HeLa cells, hiPSC-derived hNCC and undifferentiated hiPSC. Nucleolar-like foci stained by Ab H8132 are indicated by arrowheads. Ab #6364 stains endogenous NR2F1 only in the nucleoplasm and specifically in HeLa and hNCC, but not in hiPSC, which do not express NR2F1. B) Live-cell imaging of hiPSC*^NR2F1GFP^* 24 h after transfection shows nuclear but not nucleolar NR2F1 localization. Zoom-in is marked with a white box and shown for the GFP channel below. Arrowheads point to NR2F1-GFP negative nucleoli. DAPI marks nucleoplasm (blue), scale bars: 50 µm.

The confounding results of NR2F1 localization prompted us to use a system where we can avoid the use of NR2F1 Abs. Hence, we chose to transiently overexpress *NR2F1-GFP* in undifferentiated hiPSC (hiPSC*^NR2F1-GFP^*) and examined the GFP signal in living cells under a fluorescence microscope 24 h after transfection (Figure 2B). In contrast to the dominant nucleolar-like foci obtained using Ab H8132, we observed a distinct NR2F1-GFP signal only in the nucleoplasm, but not in the nucleoli, in imaging without fixation and Ab staining.

To test, whether fixation can change the NR2F1-GFP localization and to better distinguish between nucleoplasm and nucleoli, we fixed the hiPSC*^NR2F1-GFP^*with FA using our standard IF protocol^69^, and stained the cells only with DAPI. Although this made the GFP signal somewhat weaker, we again detected GFP exclusively in the nucleoplasm but not in nucleoli (Supplementary Figure 2.C first line). Since the GFP signal was weakened by FA fixation, two different anti-GFP Abs (AbGFP) were used (anti-mouse and anti-rabbit), which further confirmed the NR2F1-GFP nucleoplasmic but not nucleolar staining (Supplementary Figure 2.C second and third lines).

The assumption that the Ab H8132 signal does not reflect the actual localization of NR2F1 but rather represents a technical artefact was further supported by our bioinformatic analysis of publicly available datasets and algorithms predicting biophysical and functional domains of NR2F1. Nuclear receptors typically contain a highly disordered domain at the N-terminus^89^ and accordingly, residues 1–81 of NR2F1 are annotated as disordered (Uniprot ID: P10589). Importantly, such disordered domains have overlapping biophysical features with low complexity regions (LCRs), which have also been associated with nucleolar proteins. Low-complexity regions are typically enriched for certain amino acids, such as lysine, which we do not observe in the N-terminus of NR2F1^90–92^ (Supplementary Figure 3A). In line with the annotation as disordered in Uniprot, we saw some level of disorder in the N-terminal domain of NR2F1 as inferred by the biophysical tool IUPRED (https://iupred3.elte.hu/), which calculates the free energy of possible intra-chain interactions (Supplementary Figure 3.B). In the self-similarity dot plot only some low complexity regions were evident (Supplementary Figure 3C). Therefore, the N-terminus of NR2F1 can be describe to some degree as a low complexity region, however without the typical sequence composition. Since many nucleolar proteins are themselves direct RNA-binding proteins^93^, we also wanted to specifically probe this characteristic using DRNApred, that – among other factors – takes into account the physical properties of the amino acids as well as their predicted level of disorder^94^. As expected, an enrichment of DNA-binding residues was predicted in the annotated DNA-binding domain of NR2F1, but not a single RNA-binding residue was predicted in the protein sequence (Supplementary Figure 3D). The dissimilarity of NR2F1 to typical nucleolar proteins was further underlined by the limited overlap of described nucleolar proteins^83^ with proteins similar to the disordered domain of NR2F1 (1-81) as identified by BLASTP (Supplementary Figure 3E, white and blue circles). Only five of the 1308 proteins described as nucleolar showed local sequence similarity to NR2F1 1-81, and none of these displayed a local sequence identity higher than 61.5%. According to the supplier’s datasheet, Ab H8132 binds to the disordered domain of NR2F1. Although the nucleolar protein ZFHX4 has the highest local sequence identity with NR2F1 1-81 (61.5%), it is highly unlikely to be bound by Ab H8132 as it is not expressed in hESC^39^. Therefore, it cannot account for the nucleolar-like foci detected in hiPSC (Supplementary Figure 3.E). Lastly, we wanted to exclude the possibility that the nucleolar NR2F1 signal was due to indirect NR2F1 binding to a nucleolar protein. While this would not be able to account for nucleolar NR2F1 signal in cells that lack any NR2F1 expression, we still wanted to assess this possibility for NR2F1-expressing cells, such as hNCC, in which Ab H8132 also stained nucleoli. We therefore manually collected described NR2F1 interactors from four databases (SPRING, IntAct, BioGRID, MINT) and checked their overlap with nucleolar proteins (Supplementary Figure 3.E, white and grey circles). Five proteins were identified in the overlap, so we next evaluated their co-expression in the single-cell RNA-seq (scRNA-seq) data from hiPSC-derived NCC from^95^ (Supplementary Figure 3.F). Three of the five possible NR2F1 interacting proteins with nucleolar localization were not expressed in hNCC (BCL11B, MPPED2, C1orf210). PFDN1 and ELAVL1 were widely expressed, however the portion of cells co-expressing NR2F1 and either of both genes is 66%, with NR2F1 itself being expressed in only 68% of hNCC. Since we observed nucleolar signal in virtually all hNCC stained with Ab H8132 (Figure 1 and Supplementary Figure 1), interaction of NR2F1 with any of these nucleolar proteins cannot account for this observation, further supporting the notion, that the nucleolar dots cannot be attributed to staining of NR2F1. Thus, our results strongly suggest that the unspecific nucleolar staining in the form of foci caused by IF staining with Ab H8132 may reflect a technical artefact. However, such foci stained with this Ab have not only been reported but also quantified in several studies^60–62, 96, 97^, which may obscure some conclusions. To avoid such complications as much as possible in the future, we systematically tested Ab H8132 and Ab 6364 together with five additional antibodies in several commonly used assays and in different cell types.

### All seven Abs are capable of detecting overexpressed NR2F1 in IF, albeit to varying degrees

For our comparative study we examined a total of seven anti-NR2F1 Abs, each numbered from Ab1 to Ab7 for simplicity (Supplementary Figure 4A). The previously used Ab H8132 (hereafter referred to as Ab3) and Ab 6364 (hereafter referred to as Ab5) were also included in the panel. The distinct regions of the NR2F1 protein that have been used as immunogens to generate these Abs, are schematized in Supplementary Figure 4A and the main additional information extracted from available data sheets is summarized in Supplementary Figure 4B. To first test the Abs in IF, untransfected and NR2F1- overexpressing HEK293 cells (HEK293*^NR2F1^*) were fixed with paraformaldehyde (4% PFA for 15 minutes) 48 hours after transfection (Figure 3). In untransfected HEK293 cells, all Abs generally showed no or weak background signals (Ab6 and Ab7 weak cytoplasmic background, Figure 3A last two lines). Notably, the nucleolar-like foci were observed exclusively in the Ab3 staining (Figure 3A, third row with empty arrowheads), which is consistent with our previous results from hiPSC, hiPSC-derived hNCC, and HeLa cells (Figure 1-2). However, in HEK293 cells, the foci were much weaker, which might be related to the morphological diversity (e.g., size and number) and/or composition of nucleoli across different cells and conditions such as cell cycle or stemness^73, 98, 99^ https://www.proteinatlas.org/humanproteome/subcellular/nucleoli). Overall, all antibodies were able to detect the overexpressed NR2F1 in the nucleus, but with different intensities depending on the Ab (Figure 3B). Ab5, 6 and 7 detected NR2F1, although they were not recommended for IF according to the manufacturer (Supplementary Figure 4B). To better quantify the specificity and sensitivity of the NR2F1 Ab panel, we calculated the mean fluorescence intensities (MFI), and signal-to-noise ratios (intensity signal in transfected cells versus background in untransfected cells) of all Abs (Figure 3C-D). However, it should be noted that we used our standard IF protocol^53, 68^ for all Abs and did not optimize it for each individual Ab, for example, in terms of incubation time or other parameters that may be further adjusted. Nevertheless, the data show that using this standard IF protocol. Ab1 and Ab4 had the best signal-to-noise ratios, followed by Ab2, 3 and 5, while Ab6 and Ab7 had a rather poor ratio (Figure 3D). Importantly, regardless of the signal-to-noise ratio, Ab3 was the only antibody that additionally stained the nucleolar-like foci, but these were visible in only a few cells, apparently only those that had not been successfully transfected with the NR2F1 plasmid (Figure 3B indicated by empty arrowheads). Overall, our data strongly suggest that in the case of Ab3, the unspecific nucleolar-like staining inversely dependents on the NR2F1 protein levels. Thus, we speculate that a lower affinity and most likely unspecific target is only detected if the Ab3 cannot bind to the higher affinity target NR2F1.

**Figure 3:**
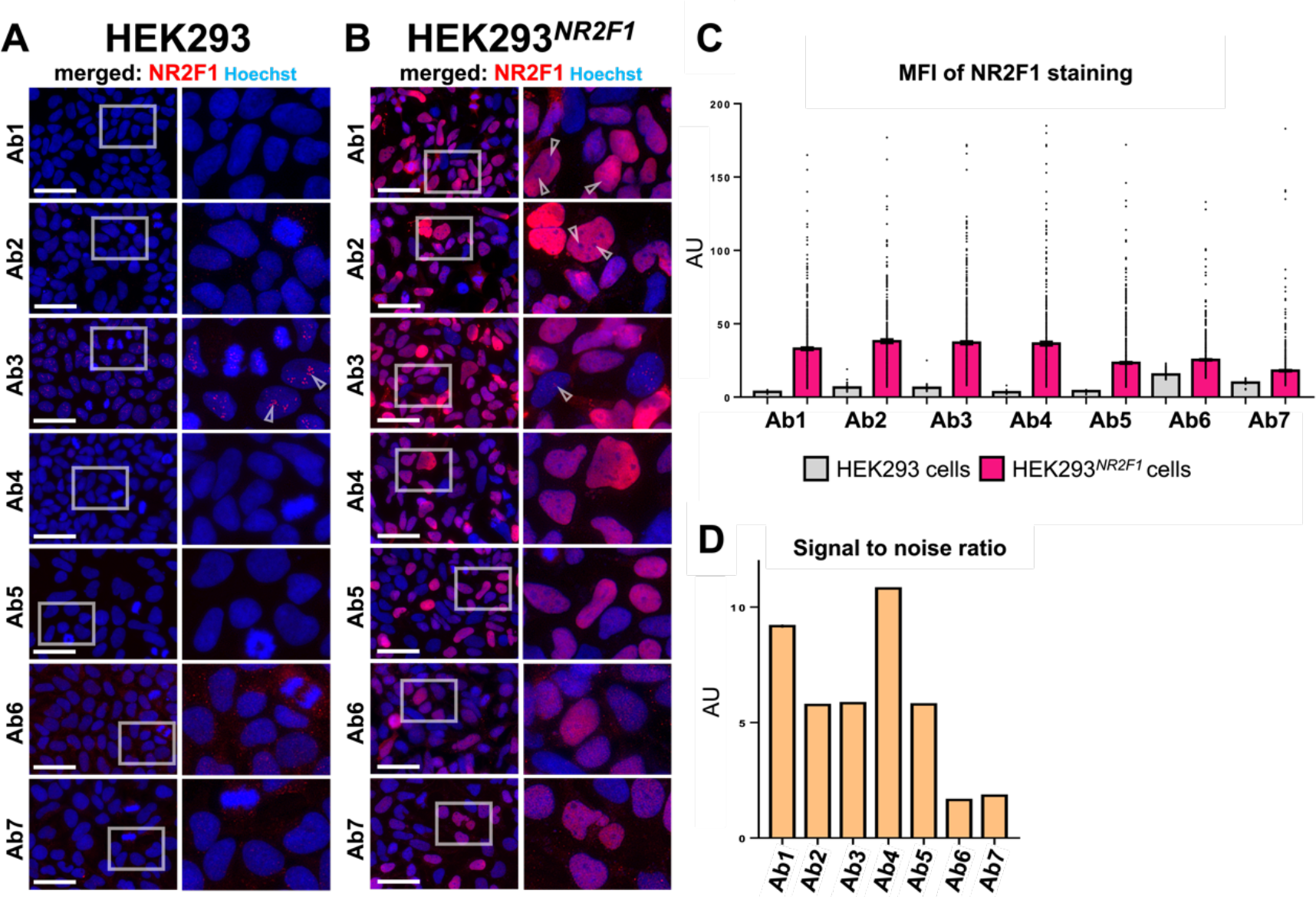
Localization and quantification of NR2F1 levels by IF in untransfected and HEK293*^NR2F1^*. A-B) The anti-NR2F1 Abs used in the IF are indicated on the left. White boxes in the first column show the selected zoom-in area placed next to it. Nuclei (blue) were stained with Hoechst. Scale bars: 50 µm. A) Negative control: untransfected HEK293 cells that do not express NR2F1 endogenously. Arrowheads indicate nucleolar-like foci exclusively in the Ab3 staining. B) HEK293*^NR2F1^*cells 48 h after transfection with a NR2F1 plasmid. Arrowheads indicate examples of nucleoli typically not stained by Ab1 and Ab2 and nucleolar-like foci stained by Ab3, seen only in cells that were unlikely to have been successfully transfected. C) MFI of histograms resulting from NR2F1 staining measured for each Ab used (indicated on the x-axis) in untransfected control HEK293 cells (grey columns) and in HEK293*^NR2F1^* (pink columns). The data are shown in arbitrary units (AU) and represented as mean ± SEM; the MFI was measured in at least 330 and up to 800 cells per sample. D) Signal-to-noise ratios between the MFI of HEK293*^NR2F1^* and untransfected HEK293 cells shown in AU as mean ± SEM for each Ab.

To corroborate the unspecific staining of NFR2F1 by Ab3, we co-stained Ab3 and Ab1 as well as Ab2 and Ab3 in untransfected (Supplementary Figure 5A) and HEK293*^NR2F1^* (Supplementary Figure 5B). Moreover, to exclude the possibility of any background signal being enhanced by the 2°Ab, we used two 2°Abs conjugated to either AlexaFlour 555 (AF555) or 647 (AF647) in two different combinations (Supplementary Figure 5A-B). And again, only Ab3 stained nucleoli-like foci in untransfected cells, regardless of the 2°Ab used, while Ab2 and Ab1 showed no significant background or unspecific staining. (Supplementary Figure 5A). In HEK293*^NR2F1^*, the NR2F1 staining resulting from Ab3 co- localized with Ab1 and Ab2 within the nucleus (Supplementary Figure 5B). In a few cases, nucleolar-like foci were observed only for Ab3, where the cells were not co-stained with Ab1 or Ab2, and therefore only in cells that were not successfully transfected with NR2F1 (arrowhead in the Supplementary Figure 5B). To further prove that the nucleolar-like foci resulting from Ab3 staining are unspecific, we stained with either Ab3 or Ab1 together with an AbGFP in hiPSC*^NR2F1-GFP^*(Supplementary Figure 5C and D). Double-positive cells for AbGFP and Ab3 staining (regardless of AbGFP or GFP signals) showed signals exclusively in the nucleoplasm, and nucleolar-like foci were visible only in GFP-negative hiPSC (Supplementary Figure 5C). In contrast, all double-positive cells for GFP and Ab1 co-localized exclusively in the nucleoplasm of successfully transfected HEK293 cells with Ab1 signal being absent in GFP-negative cells (Supplementary Figure 5D). Thus, our data provide further evidence that the nucleolar-like foci appear only when using Ab3 and are unspecific. They are particularly evident when NR2F1 expression is below a certain level that remains to be determined.

### Only four out of seven Abs are able to detect endogenous NR2F1 in hiPSC-derived NP/N, although to different degrees

Next, we systematically tested all seven Abs in IF under more physiological conditions in cells endogenously expressing NR2F1 using hiPSC-derived NP/N by a dual SMAD inhibition approach^100, 101^. Studying NP/N is of particular interest since *NR2F1* deletions or mutations result in BBSOAS, a neurodevelopmental disorder^4–6^. We compared undifferentiated hiPSC that do not express NR2F1 (Figure 2A and Supplementary Figure 2.A-B) with NP/N strongly expressing NR2F1^100^ after fixation with PFA for 15 minutes and direct stained with all Abs (Figure 4A-B). While Ab1, 2, 3, and 6 gave rise to unspecific signals in undifferentiated hiPSC, but to varying degrees and mostly in the cytoplasm, only Ab3 displayed pronounced nucleolar-like foci (Figure 4A), which had been also observed previously after FA fixation (Figure 2A and Supplementary Figure 2.B). To monitor differentiation, in addition to NR2F1, we co-stained TUJ1 to confirm the neuronal identity of the NP/N (Figure 4B-C). Overall, Ab5, 6, and 7 failed to detect endogenous NR2F1, and only Ab1, 2, 3, and 4 successfully stained endogenous NR2F1 in the nucleus, but to varying degrees (Figure 4B). Remarkably, we found that in some cells, endogenous NR2F1 was not stained or only faintly stained (Figure 4B), which was presumably due to the heterogeneity of the differentiated cells (see *NR2F1* expression in https://cancell.medisin.uio.no/scrna/hescneurodiff/ and^102^). Most importantly, and in contrast to hNCC (Figure 1), Ab3 only occasionally stained few and small nucleolar-like foci in our NP/N (Figure 4B) implying that this artefact may be cell type specific resulting from the dynamic nature of nucleoli, which varies between different cell types^1,103, 104^. Hence, determination of signal intensity by MFI values and signal-to-noise ratios in undifferentiated *WT* hiPSC and NP/N indicate that Abs1-4 are able to detect endogenous NR2F1 (Figure 4D-E). Of note, of the Abs5-7 that failed to detect endogenous NR2F1 in *WT* hiPSC-derived NP/N, Ab5 and Ab7 are not recommended for IF by the manufacturer, whereas Ab6 is (Figure 3B). To further validate our results, we used CRISPR-Cas9- engineered hiPSC in which protein expression of NR2F1 is abolished (as confirmed by WB using several Abs; Figure 7F) due to a homozygous indel in exon 2 (transcript ID: ENST00000615873.2 cut location hg38, chr5:93,585,331) that results in a premature stop codon (hereafter referred to as hiPSC*^-/-^*). Similar to *WT* hiPSC (Figure 4A-B, D-E), hiPSC*^-/-^* were differentiated into NP/N^-/-^ (Figure 4C second column) and together with undifferentiated hiPSC*^-/-^* (Figure 4C first column) all cells were stained with the seven NR2F1 Abs. The differentiated cells were additionally stained for TUJ1, a neuronal marker present in all samples, demonstrating the neuronal identity of the NP/N*^-/-^*(Figure 4C second column). In both undifferentiated hiPSC^-/-^ and differentiated NP/N^-/-^, NR2F1 antibodies Ab2, 3, and 6 showed unspecific staining as indicated by the MFI (Figure 4F). Since Ab3 still shows nucleolar-like foci in hiPSC*^-/-^*, this localization can now confidently be considered unspecific (Figure 4C first column). More generally, the data indicate that the early events of neuronal differentiation do not necessarily require NR2F1, however further characterization and functional studies of NP/N*^-/-^* were not performed here as this is outside the scope of this work.

**Figure 4:**
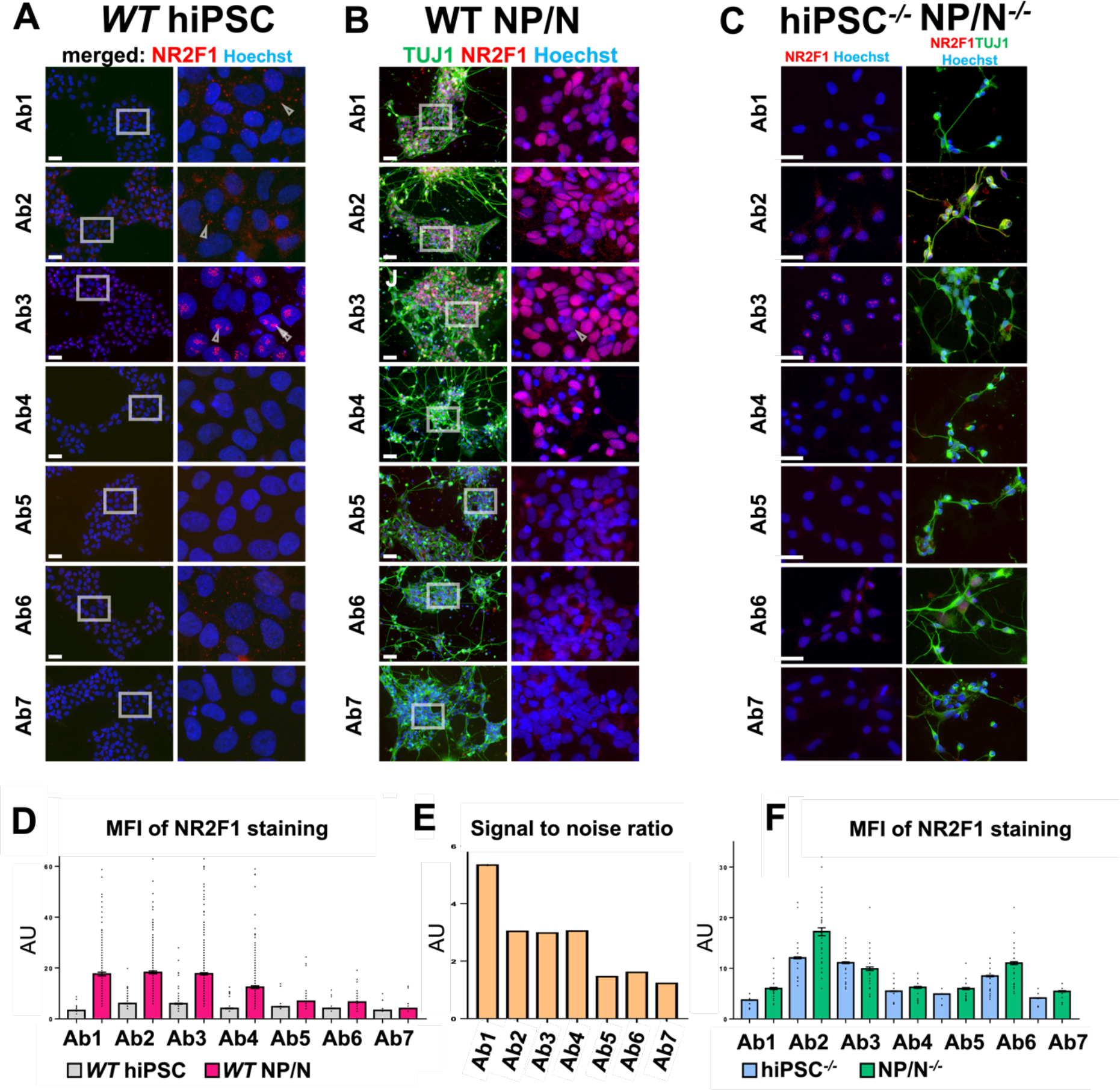
Detection of endogenous NR2F1 in *WT* and hiPSC*^-/-^*, NP/N*^-/-^* by IF using seven Abs. (A-C) Abs used to detect NR2F1 (red) are indicated on the left of each panel. The neuronal marker TUJ1 (green) was used only for IF in differentiated cells. Nuclei (blue) were stained with Hoechst. Scale bars: 50 µm. The corresponding zoom-in is next to each image. A) Undifferentiated *WT* hiPSC: arrowheads point to unspecific IF signals in the cytoplasm after Ab1 and Ab2 staining and to nucleolar-like foci after Ab3 staining. B) NP/N derived from *WT* hiPSC. C) Undifferentiated hiPSC*^-/-^* (first column) with arrowheads indicating nucleolar-like foci stained by Ab3 as well as NP/N*^-/-^*, which showed no foci (second column). D) MFI of each NR2F1 staining (Abs1-7) in AU (mean ± SEM) measured in the nucleus of undifferentiated hiPSC (grey columns) and of hiPSC-derived NP/N (magenta columns) in at least 120 cells per sample (up to 500 cells per sample). E) Signal-to-noise ratios measured for each Ab (Abs1-7) in undifferentiated *WT* hiPSC and hiPSC-derived *WT* NP/N shown in AU as mean ± SEM. F) MFI measured at least in 60 and up to 150 cells per sample of each NR2F1 staining (Ab1-Ab7) displayed as mean ± SEM in undifferentiated hiPSC*^-/-^* (blue columns) and NP/N*^-/-^* (green columns).

### IF of endogenous Nr2f1 in mouse embryonic brain sections

Since all our previous studies were based on different human cellular models, in a next step, we decided to systematically test all seven antibodies in tissue cryostat-sliced samples of mouse embryonic brain at E13.5 (Figure 5). It has been previously shown that Nr2f1 is expressed in the neocortex of the mouse brain in a gradient from anterior-low to posterior-high^4, 73^, which is essential for the formation of different cortical identities^33, 105, 106^. Hence, we compared Nr2f1 levels along the anterior-posterior axis of the developing mouse brain of *WT* and *null* mice (hereafter referred to as *KO*). This *KO* mouse model was previously generated by Cre-lox technology and the complete absence of the Nr2f1 protein has already been proved^54^. Importantly, all cryostat-sliced brain samples were subjected to a minimum of 2 h PFA fixation, which is a standard procedure for staining mouse tissue, followed by cryoprotection in sucrose and freezing^53, 68^. Figure 5A- D shows whole brain sections at E13.5 of *WT* (Figure 5A and C) and *KO* mice (Figure 5B and D) stained with either Ab1 (Figure 5A-B) or Ab3 (Figure 5C-D). In agreement with previous studies^27, 33, 107^, we observed an Nr2f1 gradient along the anterior-posterior axis of the dorsal telencephalon in the *WT* mouse, when using Ab1 or Ab3 (Figure 5A and C). In the *KO* mouse, the absence of Nr2f1 was confirmed by Ab1, but an unspecific background was observed for Ab3 (Figure 5B and D). Surprisingly, the background signal in these experimental conditions appeared to be in the cytoplasm or extracellular matrix rather than nucleoli. Therefore, to get a more accurate picture of Nr2f1 localization, we examined different cortical regions of *WT* and *KO* mice at a higher magnification for all seven Abs (Figure 5E and F). In total, only Abs1-4 were able to detect Nr2f1, but comparing *WT* anterior to posterior Nr2f1 levels, Ab1, Ab2 and Ab4 displayed the clearest gradient (Figure 5E and F). Measuring MFI and calculation of signal-to-noise ratios further confirmed our observations (Figure 5G and 6). The data suggest that the anterior-posterior Nr2f1 gradient (Figure 5C, G and H) may be masked in the Ab3 staining of the *WT* brain sections by the high background. The presence of background signals in the cytoplasm or extracellular matrix instead of nucleoli in mouse tissue stained with Ab3 suggests that the nucleoli- epitope to which Ab3 binds unspecifically may only be present in a particular cell compartment of a particular cell type. This would explain why the unspecific nucleolar-like staining is more pronounced in the nucleoli of HeLa, HEK293, hiPSC and hNCC (Figures 1 and 2, Supplementary Figures 1, 2, 4 and 4), barely visible in human NP/N (Figure 4) and absent in mouse brain tissue, but present in the cytoplasm or extracellular matrix in murine brain tissue (Figure 5). Besides cell or tissue type as covariates, the unspecific staining with Ab3 in the cytoplasm or extracellular matrix of mouse brain (Figure 5) could also be caused by the different staining protocol, namely 2 h PFA fixation, dehydration by sucrose and/or antigen unmasking by epitope retrieval eliminated during mouse tissue processing. Nevertheless, our data show that subtle changes in Nr2f1 levels in different brain regions can only be detected by the most sensitive and specific antibodies. Most importantly, the results also suggest that the cell type and/or fixation method used for sample preparation could have a major impact on the presence of artefactual nucleolar-like staining when using Ab3.

**Figure 5:**
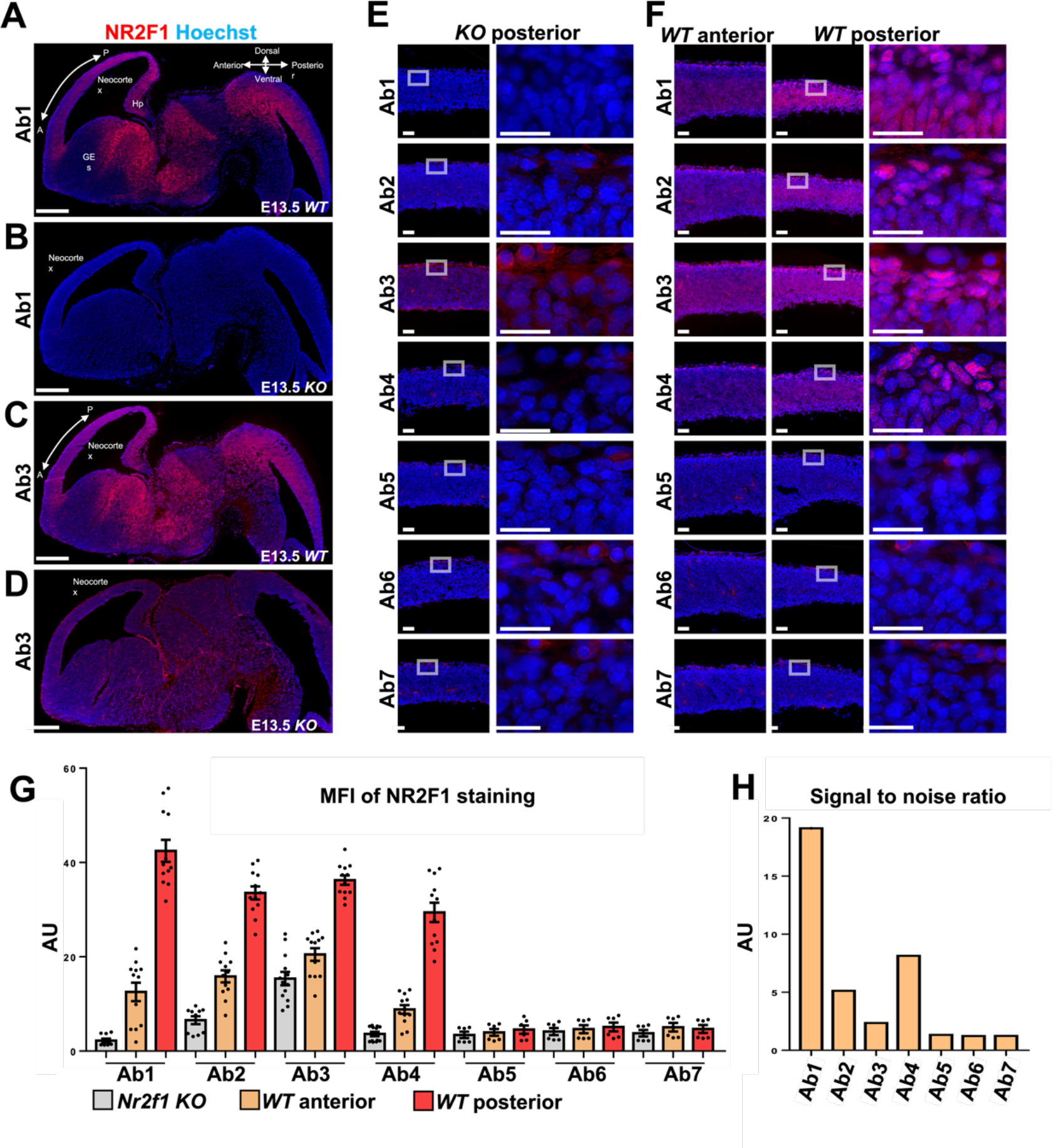
Comparison of the anterior-posterior Nr2f1 gradient in *WT* and *KO* mouse brains at E13.5 in IF using seven Abs. A-F) Sagittal sections of whole *WT* and *KO* mouse brains were stained with Abs indicated at the left side. Nuclei (blue) were stained with Hoechst. Upper right corner in A): a schematic of the anterior-posterior and dorsoventral axes. A-D) The overview of IF staining with Ab1 and Ab3 in *WT* and *KO* shows a strong background signal in the case of Ab3 in *KO*. Scale bars: 500 µm. E) Posterior images of *Nr2f1 KO*. White boxes indicate the area of zoom-in shown next to it. F) Anterior areas (first column) and posterior areas (second column) in *WT*. White boxes in E-F indicate zoom-in areas shown next to it. Only Abs1-4 can detect Nr2f1, but Ab3 leads to high background staining localized in the cytoplasm or extracellular matrix rather than in the nucleoli. Scale bars: 50 µm in first column of E) and first and second columns of F); 25 µm in last column of E) and F). G) MFI of each Nr2f1 staining measured in random fields from at least 6 and up to 12 different cortical regions per sample in *Nr2f1 KO* and *WT* brains represented as mean ± SEM in AU. H) Signal-to-noise ratios (*WT versus KO*).

**Figure 6:**
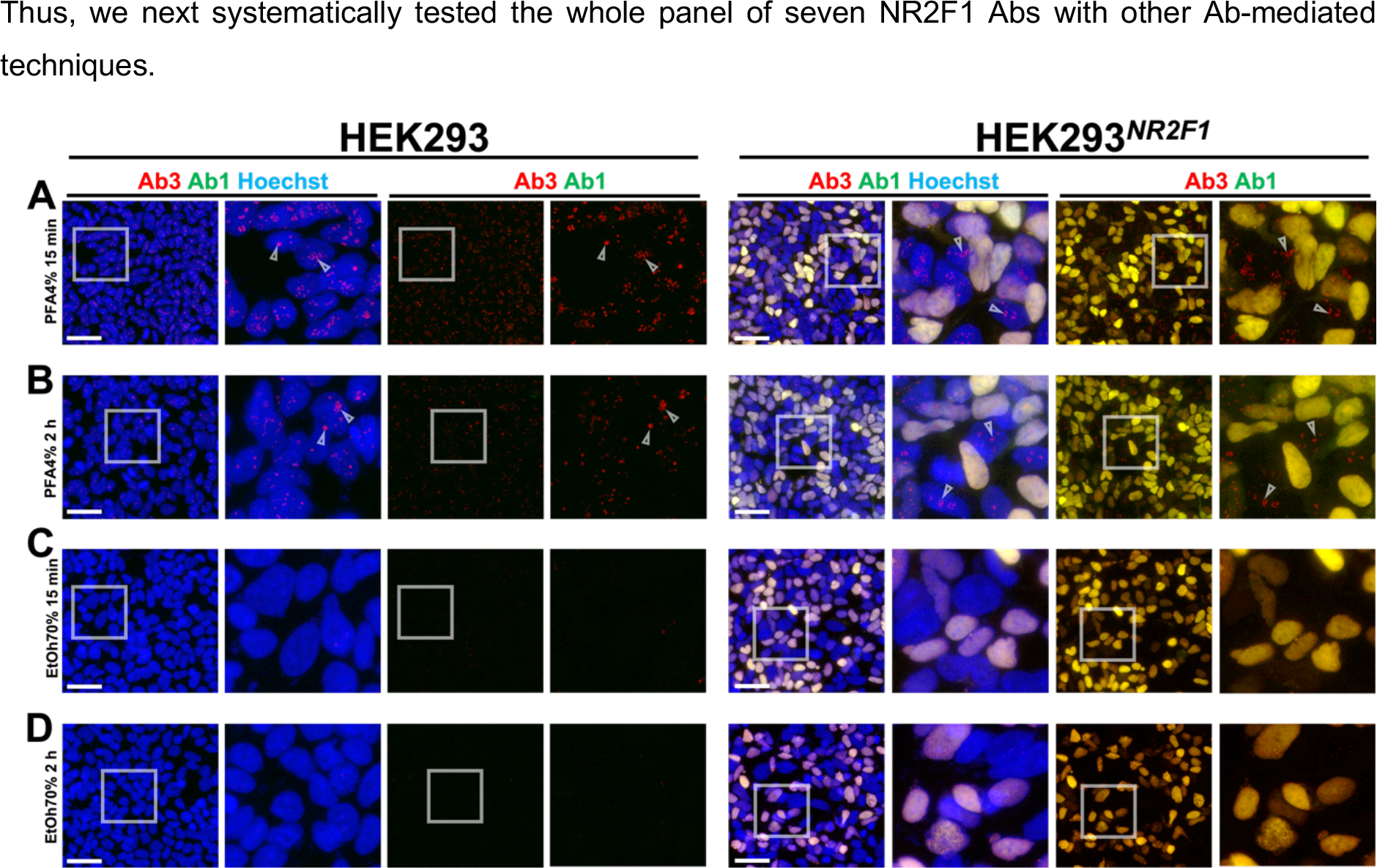
Different fixation times and conditions for IF staining with Ab3 in HEK293 and HEK293*^NR2F1^* cells. Co-staining of NR2F1 with Ab1 (green) and Ab3 (red) in untransfected cells (first four columns) and 48 h post transfection (last four columns). Only successfully transfected HEK293*^NR2F^* cells display double positive nuclei co-stained by Ab1 and Ab3. A) PFA fixation for 15 minutes and B) 2 h that reduced the unspecific nucleolar-like foci resulting from Ab3 staining without affecting the Ab1 signal. C) 15 minutes and D) 2 h fixation with 70% ethanol completely prevented the occurrence unspecific nucleolar-like foci in the Ab3 staining without affecting the Ab1 staining. Nuclei were stained by Hoechst. Scale bars: 50 µm.

**Figure 7:**
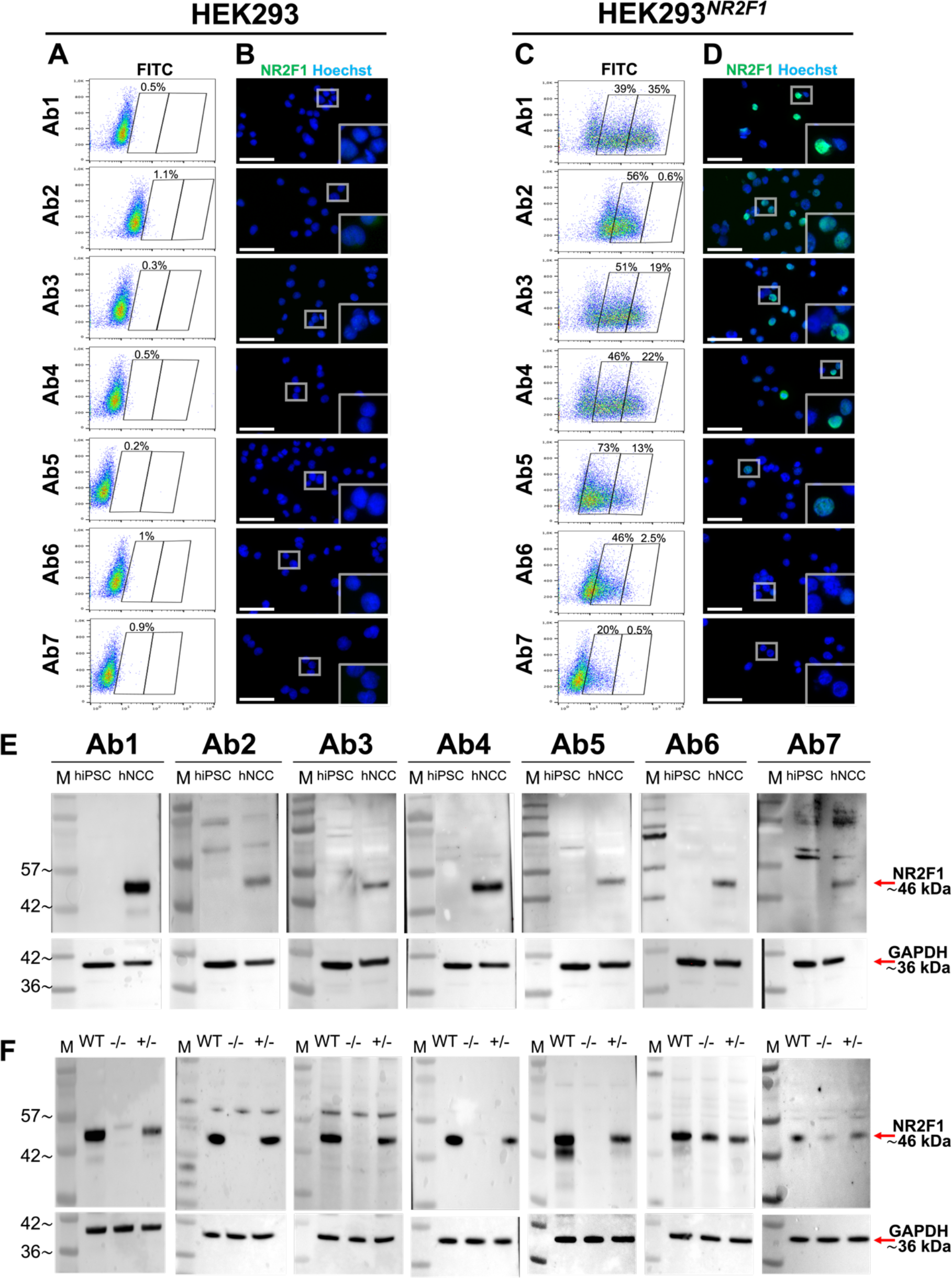
Testing all of the seven anti-NR2F1 antibodies in FC and WB. A-D) Quantification of NR2F1 levels by FC using all seven Abs in HEK293 and HEK293*^NR2F1^* cells. A-B) Untransfected and C-D) HEK293*^NR2F1^* cells were fixed with 70% ethanol 48 h after transfection, and stained with the 1°Abs indicated on the left, followed by adequate AF488 as a 2°Ab. A and C) FC data. For each sample, two batches with each 10,000 events were measured using the same settings and show that Ab1, Ab3 and Ab4 detect NR2F1 best under these conditions. B and D) After FC measurement, the cells suspensions of the remaining samples were stained with Hoechst (blue), spotted on glass slides and visualized by fluorescence microscopy. Scale bars: 50 µm. E-F) Detection of NR2F1 in WB in samples from *WT* hiPSC and hNCC as well as and hNCC*^-/-^* and hNCC*^+/-^*. 20 µg per sample of RIPA lysate was loaded. From left to right above each WB image, Abs1-7 used for NR2F1 detection are indicated. Red arrows mark a ∼46 kDa band corresponding to the specific size of NR2F1 or to GAPDH (∼36 kDa) used as loading control. E) Undifferentiated *WT* hiPSC and hNCC. F) hNCC*^-/-^* and hNCC*^+/-^*. All Abs recognized a ∼46 kDa band corresponding to NR2F1 hNCC*^+/-^*, and, depending on the Ab, several other bands. Ab6 and Ab7 also recognized a ∼46 kDa band in hNCC*^-/-^*, which was not detected by other Abs. M: Molecular size marker.

### The nucleolar-like foci observed in IF by Ab3 depend on fixation time and type

To test our hypothesis that the presence or absence of nucleolar-like foci appearing after staining with Ab3 might depend on the fixation time and/or method, we varied these parameters in the next experiment. To this end, we used HeLa cells expressing endogenous NR2F1 and compared Ab3 staining after three different fixation methods: standard cross-linking with 3.7% FA, dehydration fixation with either acetone/methanol mix or ethanol with glacial acetic acid (GA) and different fixation times (5, 10, 30 and 60 minutes). In addition, we also pre-treated the cells for 30 seconds with or without 0.5% Triton to pre-clear the nuclei (Supplementary Figure 6). Longer duration (60 minutes) of standard cross-linking with 3.7% FA with or without 0.5% Triton pre-treatment reduced not only the unspecific nucleolar-like but also the nucleoplasmic NR2F1 staining (Supplemental Figures 5A and D). In contrast, both methods of dehydration fixation without 0.5% Triton pre-treatment resulted in strong nuclear and unspecific cytoplasmic staining (supplemental Figure 5B-C). Remarkably, 0.5% Triton pre-treatment followed by 60 minutes dehydration fixation with acetone/methanol resulted in nucleoplasmic rather than focal-like nucleolar-like staining (Supplemental Figures 5E and H). A similar tendency was observed after pre-treatment with 0.5% Triton for 30 seconds (Supplemental Figures 5E) and 60 minutes dehydration fixation with ethanol/GA (Supplemental Figure 5F). Overall, our results suggest that the nucleolar-like foci stained by Ab3 can be reduced in HeLa cells endogenously expressing NR2F1 by increasing the fixation time and performing dehydration fixation instead of cross-linking fixation. (Supplemental Figure 5). These data are consistent with our results from mouse brains after IF using Ab3, in which prolonged fixation and dehydration resulted in loss of nucleolar foci but also unspecific staining of the extracellular matrix (Figure 5).

As we previously showed that the background signal and nucleolar-like appearance in IF when staining with Ab3 also depends on the NR2F1 expression level (Figure 4, Supplementary Figures 4 and 5), we next tested how the fixation protocol can influence Ab3 NR2F1 staining in our previously established HEK293*^NR2F1^* cells (Figure 3). The advantage of this system is that after transfection we obtained a population of cells with different NR2F1 expression levels. The transfected cells with high NR2F1 expression can be distinguished from non-transfected cells that do not express NR2F1 and serve as an internal negative control. Thus, we fixed HEK293*^NR2F1^* cells 48 h after *NR2F1* transfection and fixed the cells with two different fixatives for different times prior to staining with Ab1 and Ab3. Our standard 15-minute PFA fixation served as a control (Figure 6A-B). In untransfected cells, the unspecific nucleolar-like foci seen for the staining with Ab3 became partially weaker after 2 h of fixation with PFA, whereas in successfully transfected HEK293*^NR2F^* cells, nuclear NR2F1 co-stained with Ab1 and Ab3 was observed (Figure 6A-B). Therefore, we concluded that a PFA fixation prolonged to 2 h instead of 15 min leads to a slight loss of the nucleolar-like foci without increasing the unspecific cytosolic or extracellular signal (Figure 5). Finally, we fixed the untransfected and HEK293*^NR2F1^* cells with ethanol for 15 minutes or 2 h and stained them with Ab1 and Ab3. We were surprised to observe a total absence of the nucleolar-like signal in untransfected cells and only the nucleoplasmic localization of NR2F1 co-stained by Ab1 and Ab3 in transfected cells (Figure 6C-D). Taken together, our systematic tests allowed us to determine 70% ethanol as the optimal fixation method for Ab3 to obtain valid NR2F1 IF staining.

Thus, we next systematically tested the whole panel of seven NR2F1 Abs with other Ab-mediated techniques.

### Comparative analyses of seven anti-NR2F1 antibodies revealed which of them best fit for FC and WB

As our previous data strongly suggested that in some cell types the unspecific nucleolar-like staining caused by Ab3 was dependent on the level of NR2F1 expression (Figure 2, 3, Supplementary Figure 2C and 4), we used FC to quantify NR2F1 protein expression at the single cell level. To obtain a population of cells with different NR2F1 expression levels, we utilized previously tested HEK293*^NR2F1^*(Figure 3 and 6) and assessed the ability of all seven Abs to detect intracellular NR2F1 after fixation and permeabilization with 70% ethanol. Using this system, we could not only determine NR2F1- negative and -positive (NR2F1^total^) cells, but also distinguish between low (NR2F1^low^) and high (NR2F1^high^) positive cells. Furthermore, since the HEK293*^NR2F^* cells for each Ab staining were derived from the same pool of cells harvested 48 hours after transfection, it can be assumed that the same number of transfected cells were present in each sample being compared. Importantly, all samples were measured at the same voltage, which directly corresponds to the fluorescence intensity of the highly variable background of each Ab staining in the untransfected cells (Figure 7A). Overall, Ab1, Ab3, Ab4, and Ab5 showed the highest percentage of NR2F1^total^ cells (68.5-86.1%), whereby only 13.6-34.7% of the cells exhibited NR2F1^high^ levels. Although Ab5 displayed the highest number of NR2F1^total^ cells (86.1%), this corresponded mainly to NR2F1^low^ cells (72.5%), suggesting a limited ability of this Ab to detect a genuine NR2F1^high^ level upon overexpression. While Ab2 and Ab6 gave rise to only NR2F1^low^ cells (45.5-55.7%), Ab2 showed by far the lowest NR2F1^low^ signal, namely 0.54% (Figure 7C). The remaining unstained and stained cells were then plated on glass for direct visualization with a fluorescence microscope (Figure 7B and D). Although the staining intensity measured by FC and microscopy cannot be directly compared^108^, the microscopy images showed no nucleolar-like pattern of NR2F1 after staining with Ab3. This is consistent with our previous data, where Ab3 staining was more resistant to the appearance of the artefactual nucleolar-like pattern after ethanol fixation compared to FA or PFA fixation (Figure 6). Thus, our FC data show that Ab1, 3 and 4 are most suitable to measure NR2F1 expression in HEK293*^NR2F1^* cells after ethanol fixation that can be used to precisely quantify different NR2F1 expression degree at single cell level (Figure 7A-D). However, as only Ab1 is recommended for FC (Supplementary Figure 4B), NR2F1 staining with Ab2, 5, 6 and 7 for this purpose may require different fixation or further optimization.

In contrast to IF and FC in standard WB, the proteins are denatured by SDS and therefore epitopes can be recognized differently by the same Ab. An additional advantage of WB is the separation of the protein mixture by SDS-PAGE, which allows the determination of proteins according to their molecular mass and a general evaluation of Ab specificity. Hence, we systematically tested all seven Abs in WB, using *WT* hiPSC-derived hNCC that endogenously express NR2F1 and undifferentiated hiPSC that do not^88^ (Supplementary Figure 1A). As shown in Figure 7E, all Abs were able to detect a clear band of the predicted molecular weight (46 kDa)^109, 110^ in hNCC, which was not present in undifferentiated hiPSC. Whether the weak additional bands of sizes other than 46 kDa present in hNCC but not in hiPSC (Ab1) may indicate post-translational modifications or degradation of NR2F1 remains to be elucidated, as this Ab has previously been shown to generate unspecific bands in mouse head tissue^109^. Bands present in both hNCC and hiPSC, especially when Ab2 and Ab7 were used (Figure 7E) can be considered unspecific. While images of GAPDH used as a loading control were acquired uniformly with 20 seconds of exposure time for all seven membranes, different exposure times were required for each NR2F1-Ab (Supplementary Figure 7A). Of note, Ponceau S staining demonstrates an efficient and comparable protein transfer from the gel onto the nitrocellulose membrane (Supplementary Figure 7B). In general, a longer exposure time, which was especially necessary for Ab7, implies its lower sensitivity. However, similar to IF (Figures 3 and 4) or FC (Figure 7A-D), this may also indicate that the WB conditions for each individual Ab could still be optimized.

The presence of unspecific bands in WB in our and previously published samples^61, 109^ prompted us to systematically corroborate the problem using hNCC derived from *WT* hiPSC, hiPSC^-/-^ previously used for NP/N*^-/-^* (Figure 4), and from cells with heterozygous *NR2F1 knock out* (hiPSC^+/-^). Again, different exposure times of WB2 were required for the NR2F1-Abs, while they were the same for the loading control GAPDH (Supplementary Figure 7A). Overall, five of the seven Abs (Abs1-5) detected a band corresponding to the size of the NR2F1 protein (46 kDa) only in the *WT* and +/-, but not in -/- samples, implying their specificity (Figure 7F). In contrast to our previous results from hiPSC that lack NR2F1 expression shown in Figure 7E, Ab6 and Ab7 detected a 46 kDa band also in hNCC^-/-^ samples.

Considering the fact that both Abs bind to the LBD of NR2F1 (Supplementary Figure 4A), which shows very high homology between NR2F2 and NR2F1 (97%)^25, 27, 56, 111^, we assume that these two Abs may recognize NR2F2 in addition to or instead of NR2F1, since both homologs are expressed in hNCC^39, 69^. This idea is further supported by the fact that membranes stained with Ab2 and Ab4, which target the N-terminal domains of NR2F1, did not show a 46 kDa band in hNCC^-/-^ (Figure 7F). Conversely, after Ab-stripping Ab2 and Ab4 and re-staining the same membranes with Ab6 and Ab7, a distinct 46 kDa band corresponding to the molecular weight of NR2F1/NR2F2 emerged in the hNCC^-/-^ (Supplementary Figure 7C). However, for ultimate confirmation of possible NR2F1/NR2F2 Ab cross-reactivity in WB, further experimental evidence is required.

Overall, and similar to the previous results shown in Figure 7E, all Abs detected one or more bands with molecular weights other than 46 kDa, albeit to varying degrees. The majority of them are also present in hiPSC^-/-^ and hNCC^-/-^ but it remains to be experimentally tested whether and if any of them or which one of these bands observed by Ab3 may correspond to the unspecific nucleolar foci visible in IF (Figure 7E; Ab1-5). Of note, known nucleolar proteins with some local sequence identity to NR2F1 1-81 as identified by BLASTP (Supplemental Figure 3) can already be excluded because of their differing molecular masses.

## DISCUSSION

Recently, it has been shown that many nucleolar proteins are also localized to other compartments, e.g., the nucleoplasm where they exert additional functions^82, 83^. As a TF, NR2F1 is typically located in the nucleus^57–59^, however, its cytoplasmic or nucleolar localization has also been previously reported even if its potential physiological function in these compartments has not been explored^45, 60–62, 64, 65, 96^. Nonetheless, nucleolar-like NR2F1 staining detected in IF by Ab3 has previously been used to quantify NR2F1 in different tumor cells^60–62, 64, 96, 97, 112^. Architecture and assembly of nucleoli formed around specific chromosomal regions are highly dynamic processes. Nucleoli formation is regulated by phase separation making them appear as liquid droplets within the nucleus^113^ - a passive process involving weak intra- and intermolecular interactions between DNA, RNA, proteins, and lipids within a dynamic scaffold that, when disrupted, can lead to rare genetic diseases^114–116^. In addition, regulation of ribosome number seems to be particularly important for early NP^117^. Given the nucleolar-like localization of NR2F1^38, 53^, we thought it might be involved in ribosome biogenesis, which has not yet been elucidated. Thus, determining a potential noncanonical role of NR2F1 in nucleolar biology was of great interest, as it might have opened new perspectives for the study and possible treatment not only of cancer but also of BBSOAS, Hirschsprung disease, Waardenburg syndrome type IV, and the development of lymphoid and cardiac cells. Intrigued by the nucleolar-like NR2F1 localization that we also observed in hiPSC-derived hNCC using Ab3, we systematically investigated this phenomenon by employing different human cell types, *NR2F1* overexpression, CRISPR/Cas9 engineered *NR2F1 knockout* cells and mouse embryonic brain tissue.

Consistent with previously reported results^45, 60–62^, we initially showed that in hiPSC-derived hNCC endogenously expressing NR2F1, the nucleolar-like foci observed by Ab3 staining clearly colocalize with the known nucleolar protein ZSCAN1^77^. However, we did not find NR2F1 in any of the available nucleolar protein databases^83, 87, 118, 119^. Because these data lack hNCC information, we assumed that the nucleolar-like NR2F1 localization might be cell type-specific, as a cell type-dependent subcellular localization of the TF ZNF554 (Zinc Finger Protein 554) has been observed previously. In RT4 and SH- SY5Y cells, ZNF554 was found exclusively in the nucleoplasm, whereas in U2OS cells it was occasionally found in nucleoli^82^. However, we detected nucleoli-like NR2F1 foci not only in hNCC and HeLa cells endogenously expressing NR2F1 but importantly, also in hiPSC and in HEK293 cells that do not express NR2F1. Moreover, we also detected these nucleoli-like NR2F1 foci in CRISPR/Cas9 engineered hiPSC*^-/-^*, raising further strong doubts about the nucleolar-like localization of NR2F1. Remarkably, our hiPSC-derived NP/N stained with Ab3 showed only sporadically and a very weak nucleoli-like pattern of NR2F1. Moreover, in *WT* and *Nr2f1-null* mouse tissue, we did not observe such foci, but instead detected another unspecific staining, presumably of the extracellular matrix. Thus, our data strongly suggest that unspecific staining such as the nucleoli-like foci depends on the cell type and/or the procedure required for IF and makes using Ab3 for IF very questionable in some experimental conditions.

To further support our results suggesting that the nucleolar-like foci observed by Ab3 staining are unspecific, we analyzed the predicted protein properties of NR2F1. Like most nuclear receptors, the N-terminal domain of NR2F1 harboring amino acids 1-81 is annotated as disordered^89^ (Uniprot ID P10589; https://iupred3.elte.hu/). Proteins involved in nucleoli formation through liquid-liquid phase separation typically harbor intrinsically disordered domains, which have partly overlapping features with low complexity regions (LCR)^120^. Both intrinsically disordered domains and LCR have been associated with nucleoli formation among many other biological processes^113^. Moreover, LCR are typically rich in glutamic acid and lysine amino acid residues, and can be mapped by in a sequence similarity dot plot to suggest nucleolar localization of the protein^91, 92^. However, despite the annotated disordered domain in the N-terminus of NR2F1 (residues 1-81), only one glutamic acid residue and no lysine residue can be found and our dot plot analysis of the NR2F1 protein sequence did not reveal a pronounced LCR, raising initial doubts about the nucleolar localization of NR2F1. Moreover, employing a prediction tool for DNA-/RNA-binding probabilities of amino acids in protein sequences, no RNA-binding residues were predicted, although a substantial fraction of nucleolar proteins is RNA-binding^121^. In addition, we found virtually no known nucleolarly localized proteins that have high protein sequence similarity with NR2F1 using BLASTP, further reinforcing that NR2F1 is unlikely to be a nucleolar protein itself, and only very few of the known nucleolar proteins are potential interactors of NR2F1. Since co-expression of NR2F1 with these nucleolar NR2F1-interacting proteins is however not substantial enough to explain the nucleolar signal in virtually every hNCC, to date we also do not have any evidence about indirect binding of NR2F1 to nucleoli.

We noticed that the size and number of nucleolar-like foci stained with Ab3, which are likely in the (more internal) dense fibrillar component/fibrillar center subcompartments of the nucleolus, rather than the (larger) granular component, varies in different cells. More precise information would be obtained by co-staining the nuclear-like foci with Ab3 and Abs, which mark the subcompartments. However, this is beyond the scope of this paper. Nonetheless, the foci appear in HeLa cells and hNCC but barely in NP/N, although all these cells endogenously express *NR2F1*. This is not surprising, since it is known that the composition, size, morphology and number of nucleoli is dynamic and differs between embryonic and differentiated cells^1, 103, 104^. This observation supports our assumption that Ab3 recognizes a protein that is actually present in the nucleoli of some cell types only, but this would need further testing. In contrast, for other applications, such as, ChIP-seq, the detection of such nucleolar-like structure with Ab3 does not seem to interfere as reads would generally not be mapped to ribosomal genes. The underlying ribosomal sequences are highly repetitive and therefore the vast majority is not annotated in current genome assemblies (Genome Reference Consortium [https://www.ncbi.nlm.nih.gov/grc/])^81^.

One useful way to verify actual protein localization is its tagging. A classic example of experimental evidence for the nucleolar localization of an otherwise nuclear protein LEO1 (RNA polymerase-associated protein), was previously obtained using a GFP-labelled cell line^83, 122, 123^. In addition, GFP-tagged chromatin architecture protein HMGB1 (High Mobility Group Protein B 1, a nonhistone nucleoprotein and an extracellular inflammatory cytokine) was shown to localize to the nucleoplasm in live imaging but to nucleoli after FA fixation^124^. However, hiPSC*^NR2F1-GFP^* always showed exclusively nucleoplasmic but not nucleolar localization, as demonstrated by live imaging and confirmed by staining with two different AbGFP after FA fixation. Overall, our data provide strong evidence that NR2F1 does not play a role in nucleoli and strongly suggest that the nucleolar-like foci observed in IF may result from the binding of Ab3 to an unknown nucleolar target rather than to the NR2F1 protein. This implies that in certain cell types there may be a nucleolar protein that carries a very similar sequence to the part of the NR2F1 protein against which the Ab was generated. However, as mentioned above, we did not identify a nucleolar protein that has high protein sequence similarity. To exclude the possibility of Ab3 detecting the nucleolar protein with highest local sequence similarity (ZFHX4, 61.5% local sequence similarity), we found that this TF is typically not expressed in pluripotent stem cells such as hESC, making it unlikely to account for the Ab3 staining of nucleoli in virtually all hiPSC^39^. Nevertheless, it may be worthwhile to determine the potential non-NR2F1 target of Ab3 in nucleoli, e.g., by immunoprecipitation coupled with mass spectroscopy of hiPSC*^NR2F1-GFP^*compared with untransfected hiPSC. This experiment would not only reveal what this monoclonal Ab recognizes under denaturing or non-denaturing conditions, but could also reveal a potential protein with an amino acid sequence related to NR2F1 and possibly provide novel information about the structure and/or evolution of NR2F1. An interesting example of confounding data resulting from IF visualization by an Ab was reported for nuclear speckles, membraneless interchromatin space within the nucleus harboring RNA processing machinery and TFs but no DNA^125^. The Ab SC35 used in this study should principally recognize the splicing factor SRSF2, but the main targets of this Ab are other proteins SRRM2 and SON, and SRSF2 can be recognized to a lesser extent. Independent of revealing data misinterpretation using Ab SC35, the authors discovered novel roles of SRRM2 and SON in nuclear speckles formation.

We first assumed that the appearance of the nucleolar-like foci in *WT* hiPSC-derived hNCC could be explained by single-cell variability, as observed in U2OS cells for ZNF554^82^. Moreover, our results demonstrate that Ab3 is able to detect NR2F1 when it is highly expressed, e.g., in hiPSC*^NR2F1^* and HEK293*^NR2F1^*cells. Our previously published scRNA-seq data from hNCC^69^ confirmed that *NR2F1* expression is indeed quite heterogeneous and lower compared to e.g., TFAP2A and NR2F2, although the hNCC population is quite homogeneous. However, we also observed very distinct nucleolar-like staining in cells that do not express endogenous *NR2F1* such as untransfected HEK293 and *WT* hiPSC and, even more strikingly, in hiPSC*^-/-^*. Therefore, we can exclude single-cell variability as a reason for the nucleolar-like foci observed by Ab3 and our data provide further evidence that the nucleolar-like foci are rather unspecific and only occur when *NR2F1* expression is below a certain level that remains to be determined.

One of the reasons for artefacts that may occur in IF may be related to cell fixation, a step required to chemically preserve tissue for analysis by preventing cell destruction by proteases^2,3, 124^. Therefore, we tested Ab3 in HeLa cells that endogenously express *NR2F1* using different fixation types and times, as well as a method for pre-clearing the nuclei to reduce unspecific Ab-binding. In doing so, we were able to show that the nucleolar foci can be partially reduced under certain conditions, notably upon dehydration fixation and pre-clearing. To optimize these conditions, we tested Ab3 in HEK293*^NR2F1^* cells and found that no nucleolar-like foci appeared after fixation with 70% ethanol. As an organic solvent, ethanol can cause loss of small and soluble proteins, leading to speculation that the putative protein recognized by Ab3 in cells that do not express NR2F1 may be small and/or soluble. It has been shown that PFA fixation efficiency depends on the amino acid sequence of the target protein^120, 126, 127^ and tertiary structures^128^ and artefacts resulting from PFA fixation can lead to changes in the number, appearance, or disappearance of liquid condensates and thus to misinterpretation of experimental results^129^. Similar to our results observed using Ab3, it has been demonstrated that fixation can lead to artificial droplet-like foci in cells that otherwise have a homogeneous protein distribution within the nucleus with no visible spots in the live state^129^. We observed a partial improvement by the combination of ethanol with GA only in HeLa cells that were pre-cleared with Triton prior to the staining with Ab3. This could suggest that a soluble protein, that is located in the nucleolus and otherwise artificially stained by Ab3, might have been rinsed away. Pre-clearing with Triton is likely to preserve nuclear morphology, cell membrane integrity, and cytoplasmic structures, as shown by scanning electron microscopy of rat alveolar macrophages *in vitro*^130^. Both alcohol^131^ and FA^132^ have been recommended as good fixation reagents, but our IF results clearly show that the choice is highly dependent on the antibody and target protein.

Encouraged by the effect of fixation with ethanol and elimination of the unspecific nucleolar-like foci in IF after using Ab3, we tested all Abs under these conditions for FC. This method allows high-throughput quantification and effective exclusion of background staining, which is not possible with IF, at least not to this extent. Using a pool of HEK293*^NR2F1^* cells and a uniform fixation and staining protocol for all FC experiments, we demonstrated that, similar to IF, all Abs are in principle capable of recognizing NR2F1, although to a different extent. Taking advantage of the quantification and single-cell staining possible by FC, we clearly showed that under our conditions Abs1, 3, and 4 recognize overexpressed NR2F1, ranging from low to high levels, whereas Abs2 and 5 showed only low, and Abs6 and 7 very weak NR2F1 signals. Direct visualization of the remaining stained cells under the fluorescence microscope further supports our FC results. Similar to IF, individual optimization of FC staining for Abs2, 5, 6, and 7, which was not possible in this study, could potentially improve NR2F1 staining.

Overall, our systematic comparison of all seven NR2F1 Abs provided by different vendors and targeting different regions of NR2F1 revealed that Ab1-4 are able to detect endogenous NR2F1/Nr2f1 in IF, but to different extents. Interestingly, these Abs target the N-terminus of NR2F1/Nr2f1, implying that this region is best suited for the production of specific NR2F1/Nr2f1 Abs. This region is also one of the least conserved among nuclear receptors and least homologous to NR2F2^25, 89^. However, we only compared standard fixation and timing conditions for Ab3, so individual optimization for other Ab could improve IF staining of endogenous NR2F1/Nr2f1. Of note, under the same IF conditions, all Abs were able to detect highly overexpressed NR2F1 in HEK293*^NR2F1^* cells, albeit also to different extents. These results lead us to speculate that manufacturers may have validated these Abs in an overexpression system rather than in a more physiologically relevant system. However, according to the datasheets (supplementary Figure 4B) most Abs were tested in cells or tissues endogenously expressing *NR2F1.* Overall, our data suggest that manufacturers should ideally consider multimodal validation approaches that do not preclude scientists from performing the necessary control experiments in their specific biological context before answering biological questions with these Abs.

In our last Ab-based assay, we systematically tested all NR2F1 Abs in WB under standard denaturing conditions that can alter the epitope^133, 134^. To this end, we employed undifferentiated *WT* hiPSC and hNCC^-/-^ as well as hNCC*^+/-^*. When comparing *WT* hiPSC and hNCC, all seven antibodies specifically recognized endogenous NR2F1 in hNCC and not in hiPSC, which do not express NR2F1 (and *NR2F2*). However, besides the 46 kDa band corresponding to NR2F1, other bands with a different size were also detected by all Abs, most of which were also present in hiPSC. Surprisingly, two of the seven Abs recognized a 46 kDa band also in hNCC*^-/-^.* In vertebrates, in addition to NR2F1, its homologue NR2F2 has been identified, which shares 98% and 96% amino acid homology with NR2F1 in DBD and LBD, respectively^59^. Since their expression patterns and functions partially overlap and both TFs control the same as well as different genes^79^, the high homology poses a major challenge for antibody development and specificity. Moreover, the predicted molecular size of NR2F2 is 43.5 kDa, and expression of both TFs has been reported in hNCC but not in hiPSC^69^ or hESC^39^. Therefore, we assume that the band visible in hNCC^-/-^ corresponds to NR2F2 and not NR2F1, although further experimental evidence is necessary to conclusively prove this. The hNCC differentiation of hiPSC carrying a double homozygous *knockout* NR2F2 and *NR2F2*, if viable, would provide important information on the origin of this band. Notably, these cell lines could be further used to validate the previously described nucleolar localization of NR2F2 in breast cancer^78^. However, these questions are beyond the scope of our study and remain to be elucidated.

As both monoclonal and polyclonal Abs have their advantages and disadvantages, we compared mouse (Ab3) and rabbit monoclonal (Abs1 and 4) as well as rabbit polyclonal Abs (Abs2, 5, 6 and 7).

While monoclonal Abs can recognize and bind a specific epitope, polyclonal Abs can recognize and bind many different epitopes of a single antigen. Thus, monoclonal antibodies are usually less likely to cross-react with other proteins than polyclonal antibodies^135, 136^. However, in the case of Ab3, we observed unspecific nucleolar foci in IF and in WB additional bands not corresponding to the molecular size of NR2F1 in human cells that to not express *NR2F1*. Of note, Ab3 was the only mouse antibody we tested and the only one that produced unspecific nucleolar-like foci in IF. All other antibodies were rabbit Abs, but some of them showed unspecific bands in the WB. It therefore remains to be determined whether these issues are related to the species in which they were generated or to the immunogens used for their generation. Most importantly, the summary of our results (Table 1) can serve as a recommendation to the scientific community for the use of NR2F1 antibodies and the method by which we were able to detect NR2F1 in our assays. In particular, Ab1 and Ab4 performed best in our hands, independent of the sample type, fixation process or technique used.

**Table 1.**
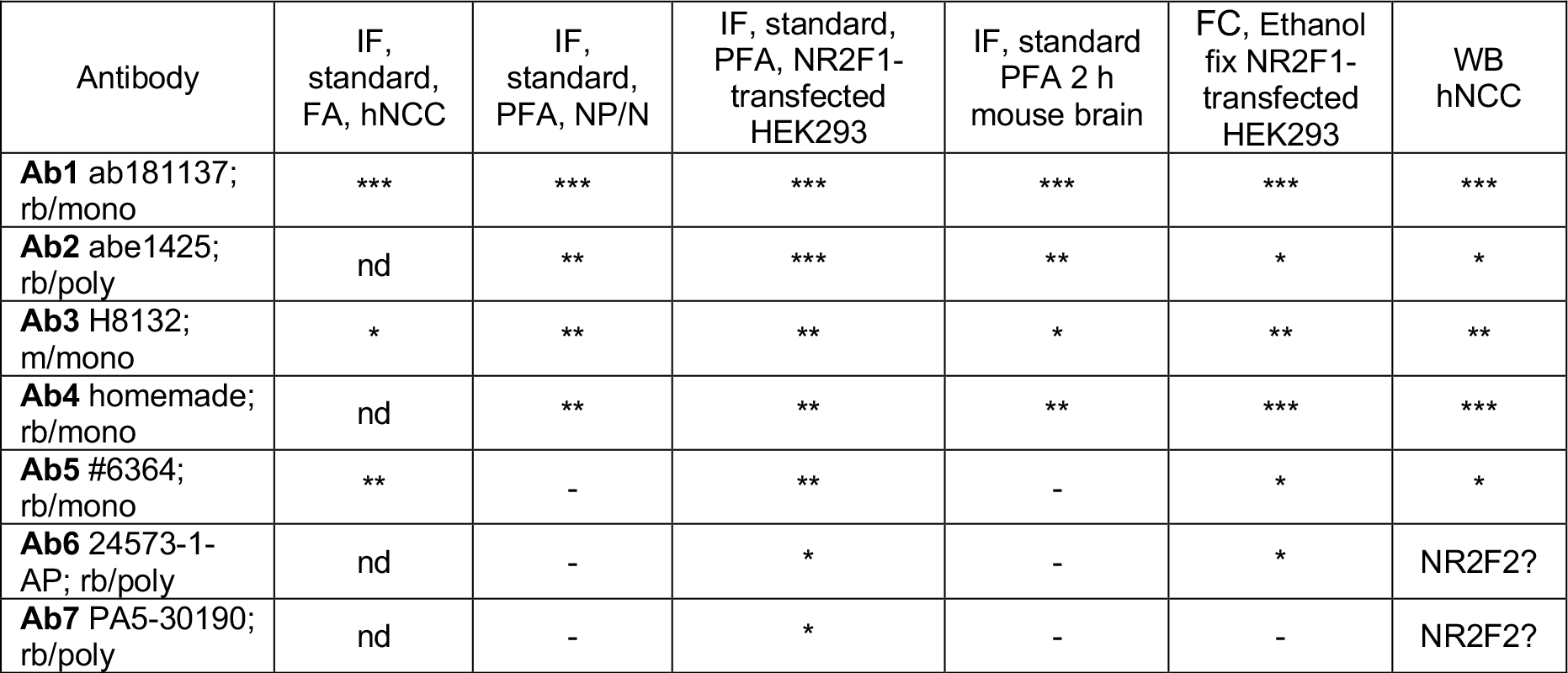
Summary of the results of the individual Abs and assays used in this study under our conditions: *** = optimal, ** = good, * = sufficient, - = no signal; nd = not determined.

There is a large number of publications on different fixation and permeabilization protocols for cells and tissues showing how important, but also how challenging it is to find the right antibodies and the right conditions^2, 3, 129, 131, 137, 138^, and our work strongly confirms these findings. Several platforms have been developed, such as antibodypedia that allows providers to post validation data in a standardized manner (https://www.encodeproject.org/, https://www.encodeproject.org/antibodies/, https://www.antibodypedia.com/), and many efforts are being made to develop it further^139^. However, this platform does not include all available Abs, and negative results are often not published. As a consequence, many scientists perform the same tests to find out that a certain antibody does not work under their conditions, which costs unnecessary time and money. Moreover, the lack of appropriate technical controls can lead to the inappropriate use of suboptimal Ab-based protocols, potentially comprising data interpretation. Therefore, it would be very helpful if scientists could share negative results using Abs in a specific context independent of the publications for which they were used. We are aware that the establishment of such a platform would need to be organized, monitored, and the data would need to be easily updated and accessible. Such a platform would be certainly be a logistical challenge, but very useful for the scientific community.

## EXPERIMENTAL MODELS; MATERIALS AND METHODS

### Cell culture

All cells were cultured under standard conditions at 37°C in a 5% CO2 atmosphere. HeLa cells were grown in DMEM high glucose (Thermo Fisher Scientific, 11965092), supplemented with 1% Penicillin-Streptomycin (Thermo Fisher Scientific, 15070063) and 10% Fetal Bovine Serum (Biochrom, 511150). Human embryonic kidney (HEK293) cells were cultured in DMEM (Thermo Fisher Scientific 41965-039) supplemented with 10% inactivated Foetal Bovine Serum (FBS; ThermoFisher 10270-106). hiPSC (DPEDi001-A) used for NR2F1-GFP overexpression and hNCC differentiation were cultured on GelTrex (Thermo Fisher Scientific, A14132-02) coated dishes in StemMACS iPS-Brew XF (Miltenyl Biotec, 130-104-368), with medium changed every other day and cells were regularly split with Versene (GIBCO by Life Technologies, 15040066.). hiPSC used for neuronal differentiation were maintained in StemFlex medium (Thermo Fisher Scientific; A3349401), on Matrigel (Corning 354234) coated cell culture plates. For routine passaging Accutase (Sigma A6964-100ML) was used and medium was supplemented with 10 µM Rock inhibitor (Y-27632; Stem Cell Technologies; #72304) for 24 h after passaging.

### Generation of HEK239*^NR2F1^* cells, hiPSC*^NR2F1-GFP^*, hiPSC^-/-^ and hiPSC^+/-^

The full-length human NR2F1 sequence was cloned into the pEGFP-C1 (Addgene-Plasmid #46956) and validated by Sanger sequencing and delivered into hiPSC using the non-liposomal FuGene HD transfection reagent (Promega, E2311) following the manufacturer’s instructions. 24 hours after transfection, cells were subjected to IF assay. To generate HEK293*^NR2F1^*, the pcDNA3.1-NR2F1-DYK (Genscript clone ID: OHu23866D) vector with the full-length human *NR2F1* sequence was transfected using JetPRIME Polyplus (POL114-07), following manufacturer’s instructions. Two days after transfections, cells were detached by Trypsin-EDTA treatment (Thermo Fisher Scientific; 25200072) and either processed for FC or seeded on Matrigel-coated round glass coverslips in 24-well plates for IF. hiPSC*^-/-^* (PGP1 c2) and hiPSC^+/-^ (PGP1 F3) were generated by Synthego using CRISPR/Cas9 technology and guide RNA sequence: CUACGGCCAAUUCACCUGCG. For PCR amplification and sequencing (Forward primer: TGGCAATGGTAGTTAGCAGCT; Reverse primer: TTGAGGCACTTCTTGAGGCG) were used and clones with an indel mutation in exon 2 of *NR2F1* causing frameshift in the DBD region and a premature stop codon which results in a truncated protein, were picked and validated by Synthego. The absence of chromosomal anomalies was verified by using a PCR-based kit (Stem Cell Technologies; hiPSC Genetic Analysis Kit, # 07550).

### hiPSC-derived hNCC

Confluent hiPSC were differentiated into hNCC as described in Laugsch et al., 2019. The colonies were detached by 2 mg/mL collagenase, washed with PBS and plated in Petri dishes in hNCC differentiation medium (Neurobasal and DMEM F12 media in 1:1 ratio, 0.5x B27 with Vitamin A and 0.5x N2 supplements, 20 ng/mL epidermal fibroblast growth factor, 20 ng/mL basic fibroblast growth factor and 5 µg/mL Insulin) that was changed every 2-3 days. The resulting embryoid bodies (EB) typically attached at day 5-6 and gave rise to NCC outgrowths, which were harvested at day 11 using Accutase (Sigma-Aldrich, A6964). 50,000 cells per cm^2^ were seeded on cell culture dishes coated with 5 mg/mL fibronectin in hNCC maintenance medium in which insulin was replaced by 1 mg/mL BSA. Following one additional passage, hNCC cells were either seeded on round glass coverslips for IF or in 6 well-plates for WB and harvested in passage number 2.

### hiPSC-derived NP/N

We used a previously reported differentiation protocol based on dual inhibition of SMAD signalling^101^, with some modifications. Neural induction medium added to hiPSC at 80-90% confluency consisted of DMEM/F12 and supplemented with: N2 (Thermo Fisher Scientific; 17502-048), B27 (without vitamin A; Thermo Fisher Scientific; 12587-010), Glutamax (Thermo Fisher Scientific 35050-038), Na-Pyruvate (Sigma S8636), Penicillin-Streptomycin (Biozol ECL-ECB3001D) or Antibiotic/Antimycotic solution (Sigma A5955-100ML), β-Mercaptoethanol (0.05 mM; Thermo Fisher Scientific 31350-010), NEAA (1 mM; Thermo Fisher Scientific 11140-035) and Heparin (2 µg/ml; Sigma H3149-25KU) and supplemented with LDN-193189 (0.25µM; Sigma SML0559-5MG) and SB-431542 (5µM; Sigma S4317-5MG). As soon as neural rosettes appeared (7-8 days after induction) the cells were cultured in the induction medium without LDN and SB for up to 1 month, by changing medium every other day. For redistributing cells prior to IF, cells were detached by Accutase (Sigma A6964- 100ML) and seeded on Matrigel-coated round glass coverslips for one week.

### Antibodies and fluorescents staining DNA

Abs and reagents are listed in table 2 including concentrations used in IF, WB and FC.

**Table 2:** List of all antibodies and fluorescents staining DNA including concentrations used in all assays in this study.

| Abs used in this study | Company | Ref number | Conc. in T12 hiPSC, hNCC and HeLa cells | Conc.in HEK293 cells, PGP1 hiPSC, NP/N and mouse brain | Stock |
| --- | --- | --- | --- | --- | --- |
| Ab1 | Abcam | ab181137-Rabbit-Monoclonal antibody | WB - 0.4 μg/ml<br>IF - 1,6 μg/ml | 0.4 μg/ml | 0.4 μg/μl |
| Ab2 | Sigma-Aldrich | abe1425-Rabbit-Polyclonal antibody | WB - 1:500 | 1:1000 | not specified |
| Ab3 | Perseus Proteomics | PP-H8132-00- Mouse-Polyclonal antibody | WB -1 μg/ml<br>IF - 4 μg/ml | 1 μg/ml | 1 μg/μl |
| Ab4 | Homemade-purified | Rabbit-Monoclonal antibody | 1:1000 | 1:1000 | not specified |
| Ab5 | Cell signaling | 6364- Rabbit-Monoclonal antibody | WB - 1:1000<br>IF - 1:1000 | 1:1000 | not specified |
| Ab6 | Proteintech | 24573-1-AP- Rabbit-Polyclonal antibody | WB - 0.6 μg/ml | 0.4 μg/ml | 0.4 μg/μl |
| Ab7 | Thermo Fisher Scientific | PA5-30190- Rabbit-Polyclonal antibody | WB - 0.92 μg/ml | 0,92 μg/ml | 0.92 μg/μl |
| GAPDH | Abcam | ab8245- Mouse-Monoclonal antibody | WB - 0.4 μg/ml | not used | 2 μg/μl |
| AbGFP Mouse | Santa Cruz | sc-9996- Mouse-Monoclonal antibody | IF - 20 μg/ml | not used | 2 μg/μl |
| AbGFP Rabbit | Thermo Fisher Scientific | A-11122-Rabbit-Polyclonal antibody | IF - 1 μg/ml | not used | 2 μg/μl |
| HRP Conjugate | BioRad | Goat Anti-Rabbit IgG (H + L)-#1706515 | WB - 1:5000 | not used | not specified |
| HRP Conjugate | BioRad | Goat Anti-Mouse IgG (H + L)- #1706516 | WB - 1:5000 | not used | not specified |
| AF488 Mouse | Thermo Fisher Scientific | # A-11001 | IF - 2 μg/ml | 2 μg/ml | 2 μg/μl |
| AF488 Rabbit | Thermo Fisher Scientific | # A-11008 | IF - 2 μg/ml | 2 μg/ml | 2 μg/μl |
| AF594 Mouse | Thermo Fisher Scientific | # A-11005 | IF - 2 μg/ml | 2 μg/ml | 2 μg/μl |
| AF594 Rabbit | Thermo Fisher Scientific | # A-11012 | IF - 2 μg/ml | 2 μg/ml | 2 μg/μl |
| TUJ1 | Sigma-Aldrich | T8660 | not used | 1 μg/ml | 1 μg/μl |
| DAPI | Thermo Fisher Scientific | # 62248 | IF - 1 μg/ml | not used | 1 μg/μl |
| Hoechst | Invitrogen | H3570 | not used | 1:10000 | 10 mg/ml |

### Immunofluorescence of human cells

As mentioned previously, all cells used for IF were cultured on round glass coverslips in medium appropriate for the cell type. hNCC, HeLa or undifferentiated *WT* hiPSC or transfected with *pEGFP-NR2F1* (hiPSC*^NR2F1-GFP^*) were fixed with 3.7% FA for 15 min, permeabilized with 0.5% Triton X-100 in PBS for 10 min and blocked with 5% BSA/PBS for 30 min at RT. The specific reagents and fixation times used for the optimization of the Ab3 staining in HeLa cells and are indicated in the results. 1°Ab, were incubated in 5% BSA/PBS over night at 4°C. Following three washing steps with PBS 0,05% Tween, 2°Abs diluted in 5% BSA/PBS were incubated for 30 min at 37°C. After three washing steps with PBS 0,05% Tween, the nuclei were stained with 1µg/ml DAPI solution in PBS for 1 min at room temperature. Subsequently, the cells were rinsed three times with PBS and permanently mounted in Fluoromount-G (Life Technologies) and analyzed with a Leica DFC 7000 T fluorescence microscope with LAS X Software, then assembled and labelled by ImageJ2 (using FiJi package) and edited by PhotoShop. HEK293 cells were fixed in 4% PFA (Sigma P6148-1KG) for 15 minutes at room temperature. The modified fixation times and reagents used for Ab3 staining optimization in HEK293 cells are indicated in the results. After fixation, cells were washed with PBS and blocked for 1 hour with 5% sheep or fetal calf serum and 0.3% Tween or Triton diluted in PBS. Following incubation of 1°Abs (4h at RT or overnight at 4°C), and three washing steps with 1% serum in PBS, 2°Abs were incubated at RT for 2-4h. Hoechst or DAPI were included in the 2°Ab solution (10,000 x dilution). Images were acquired with an Apotome Zeiss using AxioVision software and edited by PhotoShop or by ImageJ/Fiji.

## Animals

All mouse experiments were conducted in accordance with relevant national and international guidelines and regulations (European Union rules; 2010/63/UE) and have been approved by the local ethical committee in France (CIEPAL NCE/2019-548). Homozygous *Nr2f1 KO* embryos were generated and genotyped as previously described^54^. Littermates of *KO* embryos with *WT Nr2f1* alleles were used as control mice. Midday of the day of the vaginal plug was considered as embryonic day 0.5 (E0.5). Control and mutant mice were bred in a 129S2/SvPas background. Both male and female embryos were used in this study; age is specified for each embryo used in specific experiments. Standard housing conditions were approved by local ethical committee; briefly, adult mice were kept on a 12 h light-dark cycle and housed three per cage with the recommended environmental enrichment (wooden cubes, cotton pad, igloo) with food and water *ad libitum*.

## Immunofluorescence of mouse embryonic brain sections

A detailed protocol for immunostaining of cryostat sections has been described elsewhere^68^. Briefly, mouse embryonic whole heads were dissected and fixed in 4% PFA (Sigma P6148-1KG) at 4°C for 3 to 4 hours in agitation, then washed in PBS and dehydrated in 25% sucrose overnight at 4°C. After embedding in OCT resin at -80°C, 12 µm-thick slices were obtained by cryostat sectioning and collected on SuperFrost slides (Thermo Fischer Scientific, Superfrost Plus: J1800AMNZ). Following antigen retrieval prior to incubation (10 min at 95°C in pH=6 Citric acid solution) the samples were blocked with 5% serum and 0.3% Tween or Triton in PBS for 1h. The 1°Ab were incubated for 4h at RT or overnight at 4°C and after subsequent three washing steps with PBS, the samples were incubated with adequate

2°Ab for 2-4h at RT. Hoechst or DAPI were included in the 2°Ab solution (10,000 x dilution). For mounting, a solution of 80% glycerol and 2% N-propyl gallate was used. A detailed version of this protocol has been previously published on Bio-Protocol^68^. The images were acquired at an Apotome Zeiss, using the AxioVision software and edited by PhotoShop.

### Statistics

IF data was statistically analyzed and graphically represented using Microsoft Office Excel software and GraphPad Prism (version 7.00). Quantitative data are shown as the mean standard error (SEM). For mean fluorescence intensity (MFI) quantification after immunofluorescence, measurements were performed on at least 100 cells or on at least 6 brain areas, unless otherwise stated. To calculate the MFI, we used FIJI in-built function "Adjust, Threshold" to detect nuclei/cells, followed by the "Analyze particles" and "Measure" functions to automatically quantify the pixel intensity in the areas of interest. For mouse brain, Photoshop was used instead, by taking advantage of the "Magnetic Lasso" tool to manually select brain areas of interest, followed by the "Measurement" function to quantify the average fluorescence intensity inside the selected areas.

### Flow cytometry

HEK293 cells were dissociated into a single cell suspension by using trypsin-EDTA and transferred into low-bind Eppendorf tubes, washed twice with ice-cold PBS, and fixed with 70% ethanol while mixing on a vortex, and stored at -20°C. 200,000 cells were then stained in suspension using the same IF protocol as for cryostat sections, with the only difference that a short centrifugation (5 min at 170 g) was used for the washing steps with 1% serum PBS. For each staining, 10,000 total events (excluding debris) from two independent batches (20,000 cells in total) were measured at the same voltage using the FITC (488) laser with the BD LSRFortessa system and FACSDiva software, and the data were then analyzed using FlowJo software (Becton Dickinson).

### Western blot of hiPSC and hiPSC-derived hNCC

The cells were washed with PBS and incubated on ice for 1 hour with ice-cold RIPA buffer (50 mM Tris-HCL, pH 7,5 with 150 mM sodium chloride, 1.0% NP-40, 0.5% sodium deoxycholate and 0.1% sodium dodecyl sulfate) supplemented protease & phosphatase inhibitor cocktail (100x) (Thermo Fischer Scientific #1861281). Following centrifugation for 30 min at 4 °C and at 13,000 rpm, the supernatant was collected and the protein concentration was determined using Pierce™ BCA Protein Assay Kit (Thermo Fischer Scientific, 23225). Subsequently, 20µg of each sample was boiled in a Laemmli sample buffer (Bio-Rad) containing 50 mM β-mercaptoethanol. Finally, the proteins were separated by SDS-PAGE (NuPAGE 4-12%, Bis-Tris gel (Thermo Fischer Scientific, NP0322BOX) using XCell4 SureLock Mini-Cell Electrophoresis System (Thermo Fischer Scientific, EI0001) and transferred onto a nitrocellulose membrane (GE Healthcare Life science, 10600002) using the Mini-PROTEAN Tetra cell blotting device (BioRad, 1660827EDU). The membranes were blocked for 1h with 5% BSA/PBS or 5% non-fat milk/TBS at RT and incubated overnight at 4°C with 1°Ab diluted in 5% BSA/PBS or 5% non-fat milk/TBS depending on manufacture’s recommendation. After three washing steps with 0.05% Tween in PBS, the membranes were incubated for 1h at RT with HRP Conjugate 2°Ab diluted in 5% BSA/PBS. Following three washing steps with 0.05% Tween in PBS the proteins were visualized by chemiluminescence using Clarity TM Western ECL Substrate kit (BioRad, 170-5060) and an INTAS ECL ChemoStar imaging system.

### Bioinformatical analyses

DNA/RNA-binding probabilities for all residues in the NR2F1 protein sequence were calculated by the DRNApred algorithm^94^ through the web-server interface (http://biomine.cs.vcu.edu/servers/DRNApred/). Protein BLAST (BLASTP) with NR2F1 1-81 as query sequence was run on https://www.uniprot.org/blast. Gapped local sequence alignments (gap opening cost: 10, cap extension cost: 1) were performed against human proteins with a BLOSUM80 matrix. The resulting proteins (all with E-value < 0.01) corresponded to 118 unique genes. Described NR2F1- interacting proteins were collected from BioGRID, IntAct, MINT and STRING. The overlap of both BLASTP results and the list of unique proteins physically interacting with NR2F1 with previously described nucleolar proteins^83^, was visualized using the *eulerr* package (7.0.0) (https://cran.r-project.org/package=eulerr) in *R* (4.2.2) (R Core Team, 2022 [https://www.R-project.org/]). DRNApred results were visualized with the *geom_tile()* function from *ggplot2* (3.4.1). The scRNA-seq dataset from hiPSC-derived NCC was previously described^69^. For visualization of NR2F1 expression and some of its putative nucleolar interacting proteins, briefly, count data was preprocessed using standard *SingleCellExperiment* (1.20.0) workflow^140^ and *Seurat* (4.3.0)-based visualization^141^.

### Dot plot generation

Self-comparison dot plot of NR2F1 (Uniprot ID: P10589) was generated as previously published (Lee et al. 2022). Every position in the NR2F1 protein sequence was compared to every other position in the protein in a 2D-matrix, and matches were assigned foci.

## ACKNOWLEDGEMENTS

We would very much like to thank Prof. Dr. Argyris Papantonis from the University of Göttingen in Germany for his discussion and feedback on the project and the manuscript. In addition, we thank Dr. Nima Jaberi-Lashkar from Massachusetts Institute of Technology, Cambridge, United States created a dot plot and assisted us with his expertise in the biology of nucleolar proteins.

## AUTHOR CONTRIBUTUIONS

MB, ST and ML conceived the study and designed the experiments. Experimental data were generated and analyzed by MB, ST, AA, ML with the help of AL, GM, WP, KC and ME. AC analyzed the flow cytometry data and supported us with her expertise in interpreting the flow cytometry data. ME performed bioinformatic analyses of publicly available data. MB, ST, AA and ML assembled all figures. MS and CS provided cell lines, participated in the discussion of results, and edited the manuscript. ML wrote the manuscript with input from MB.

## INCLUSION AND DIVERSITY

We support inclusive, diverse, and equitable conduct of research.

---

**Supplementary Figure 1.**
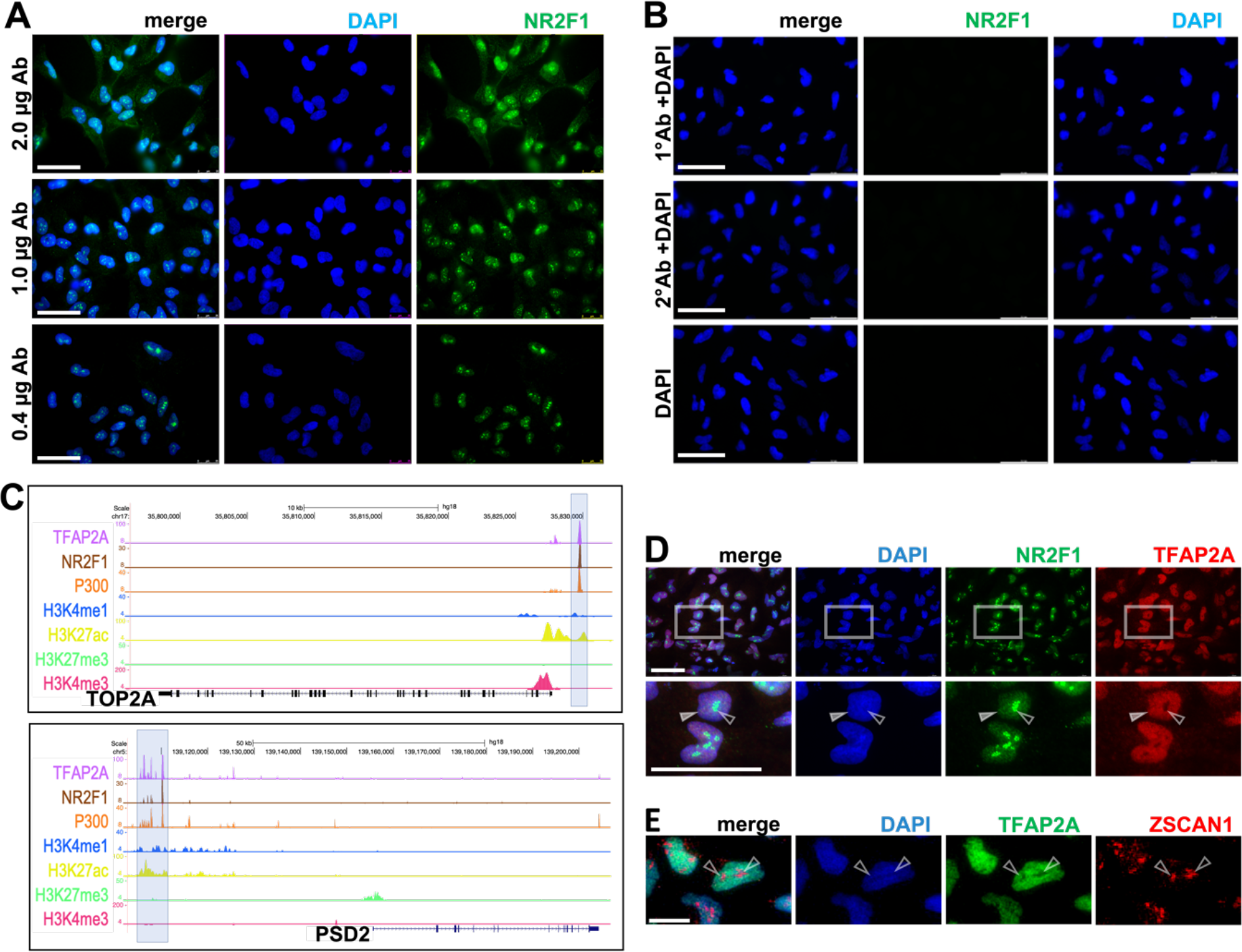
**(related to Figure 1): Validation of endogenous NR2F1 IF-staining by Ab H8132 in hiPSC-derived hNCC.** A) Testing of different Ab H8132 concentrations: 2 µg, 1 µg and 0.4 µg/ml. B) Control staining by sole 1°Ab (Ab H8132) and sole 2°Ab anti-mouse AF488. C) UCSC Genome Browser view (hg18) of publically available ChIP-seq data^39^ performed in hNCC derived from human embryonic stem cells (hESC) using Ab H8132. Examples of genomic loci (*TOP2A* on chromosome 17 and *PSD2* on chromosome 5) with active enhancer chromatin states co-occupied by TFAP2A and NR2F1 (transparent blue bars). ChIP-seq signals for H3K27ac (yellow), H3K4me1 (green), and p300 (orange) map active enhancers that are co-occupied by TFAP2A (purple) and NR2F1 (brown). D) Co-IF of NR2F1 (green) and TFAP2A (red) at low magnification (upper line) and high magnification (second line). White arrowheads indicate nuclear staining seen for both TFs; empty arrowheads highlight aggregates observed only for NR2F1. E) Co-IF of ZSCAN1 (red) with TFAP2A (green) showed no nucleolar-like co-localization. Scale bars: 20 µm.

**Supplementary Figure 2.**
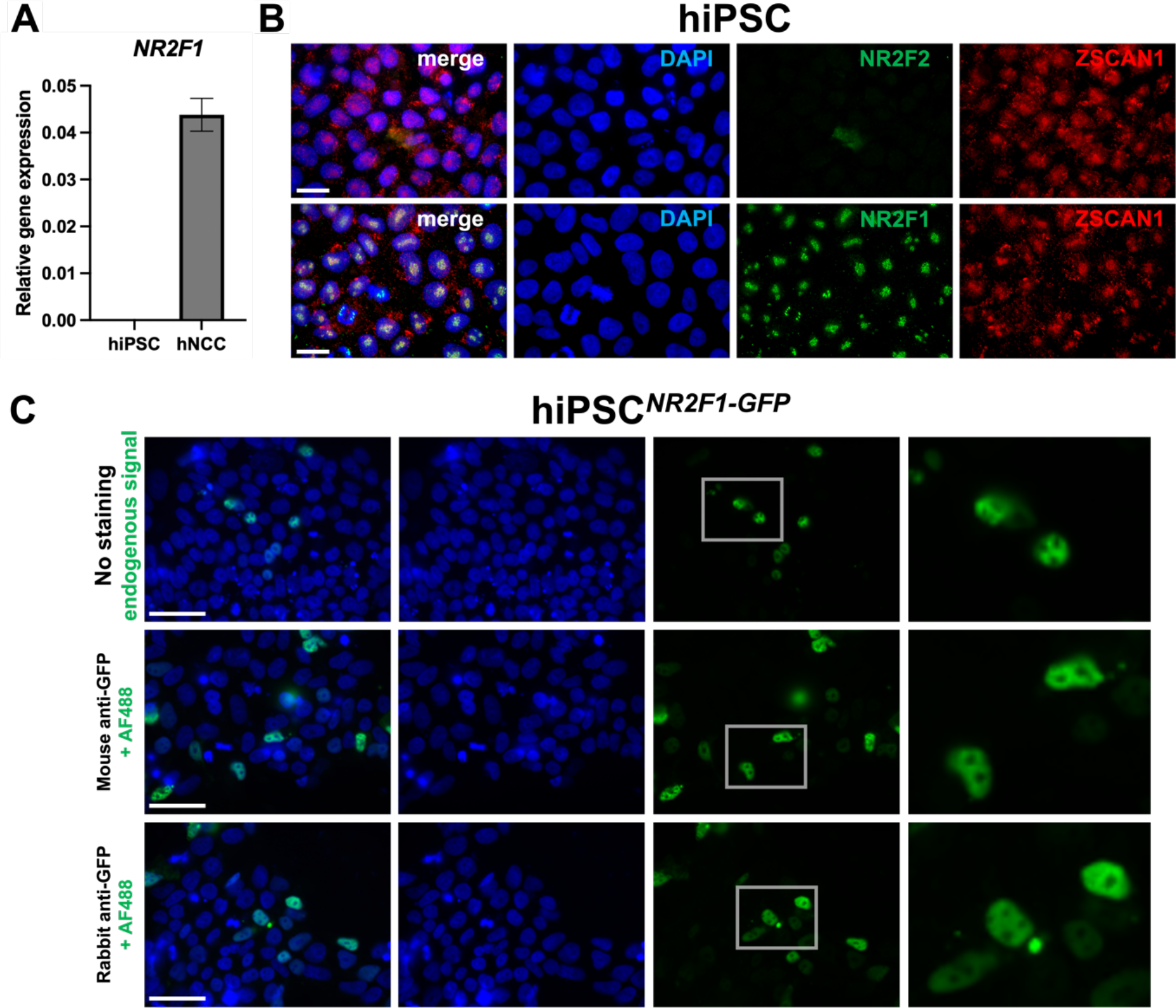
**(related to Figure 2): Localization of endogenous and overexpressed NR2F1 in undifferentiated hiPSC.** A) RT-qPCR data of the relative expression of *NR2F1* in *WT* hiPSC and hiPSC-derived hNCC showing that *NR2F1* is not detectable in hiPSC. B) Co-IF of the nucleolar marker ZSCAN1 with NR2F1 or NR2F2 in hiPSC (the latter is also not expressed in hiPSC^69^). The nucleolar marker ZSCAN1 (red) does not co-localize with NR2F2 (green) (first line), but with NR2F1 (green) when stained with by Ab H8132 (second line). C) hiPSC*^NR2F1-GFP^* (24 h post transfection) after fixation and staining with DAPI (first line) and after staining with anti-mouse or anti-rabbit AbGFP and DAPI. NR2F1-GFP is visible in the nucleoplasm without any nucleolar-like foci (second and third lines). Scale bars: 20 µm in B, 50 µm in C.

**Supplementary Figure 3.**
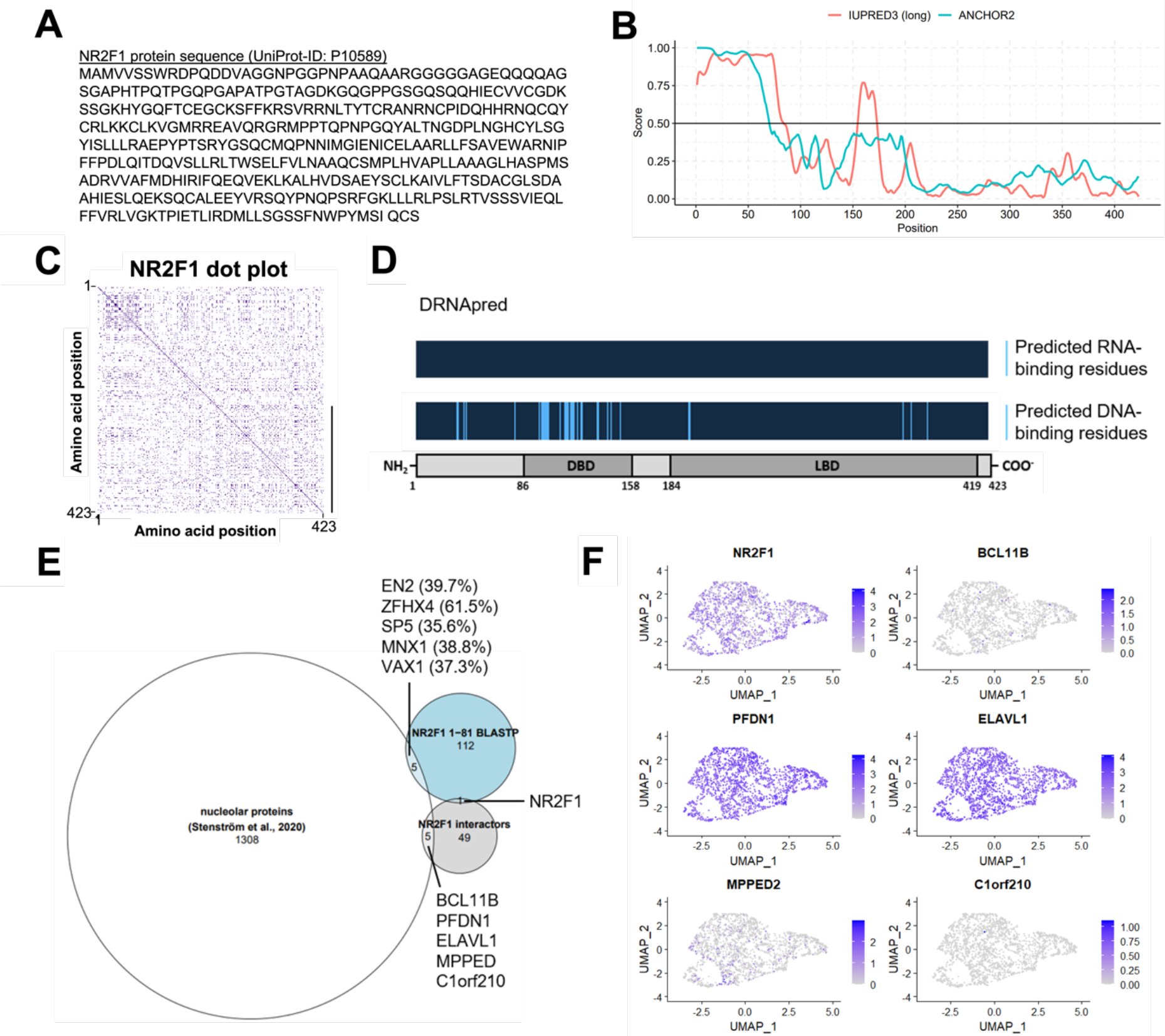
(related to Figure 2): Bioinformatical analysis of NR2F1 protein domains, putative interacting proteins and their expression in scRNA-seq. A) Amino acid sequence of the NR2F1 protein showing no enrichment of lysine or glutamic acid typical for LCRs. B) IUPRED prediction of disordered regions in the protein sequence of NR2F1. The score as a measure for the level of disorder based on a biophysical model of intra-chain interactions along the full length of the protein sequence shows some level of disorder only in the N-terminal region and the in the hinge domain between DBD and LBD. ANCHOR predicts the N-terminus as a possible disordered binding region. C) Self-comparison dot plot of the NR2F1 protein sequence. Every position in the NR2F1 protein sequence was compared to every other position in the protein in a 2D-matrix. N-terminus to C-terminus is shown from top to bottom, and left to right on the dot plot. Only very weakly dense regions appear along the diagonal in the N-terminus, indicating some LCRs. D) DRNApred predicts enrichment of DNA- binding residues in the DBD of NR2F1 and no RNA-binding residues. E) Human proteins showing sequences similar to the disordered domain of NR2F1 identified with BLASTP (blue circle). Their overlap with known nucleolar proteins is only marginal (n=5). Only five out of 1308 of known NR2F1- interacting proteins (grey circle) collected from BioGRID, IntAct, MINT, and STRING are not localized to nucleoli. F) scRNA-seq data from Laugsch et al., 2019 confirmed that only a fraction of hiPSC-derived hNCC co-express NR2F1 and any of the five potential nucleolar interactors (<68%). Protein-protein interaction with any of these proteins is therefore unlikely to account for the nucleolar Ab H8132 signal, that we observed in virtually all hNCC (Figure 2A).

**Supplementary Figure 4.**
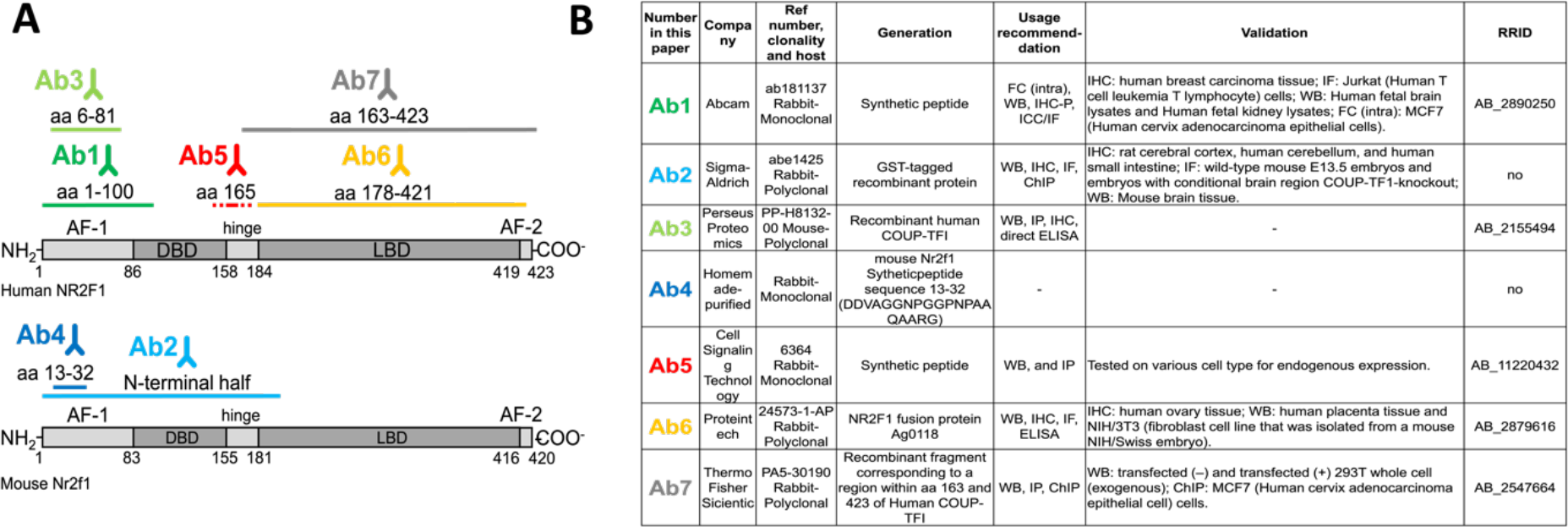
(related to Figures 3-7): Overview of immunogen regions recognized by the anti-NR2F1 Abs tested in this study and complementary information. A) Grey: schematic structure and domains of the human NR2F1 (top) and the murine Nr2f1 protein (bottom). The immunogens used to produce the Abs are indicated in different colors above the NR2F1 domains used as antigen. B) Summary of available information, including clonality, host, and immunogen. IHC: immunohistochemistry; IHC-P: immunohistochemistry, paraffin; FC (intra): flow cytometry, intracellular; ELISA: enzyme-linked immunosorbent assay, RRID: Research Resource Identifiers. Note th at all Abs are commercially available, with the exception of Ab4, which was a homemade one, produced and purified by Thermo Fischer Scientific using the mouse Nr2f1 peptide sequence 13-32 as an immunogen. Now available commercially, Ab1 has been produced as described elsewhere^55^. Importantly, the very high amino acid sequence homology between human NR2F1 and mouse Nr2f1 proteins, which share the same domains with only a few amino acids shifted, allows the use of the antibodies for both species^25^.

**Supplementary Figure 5.**
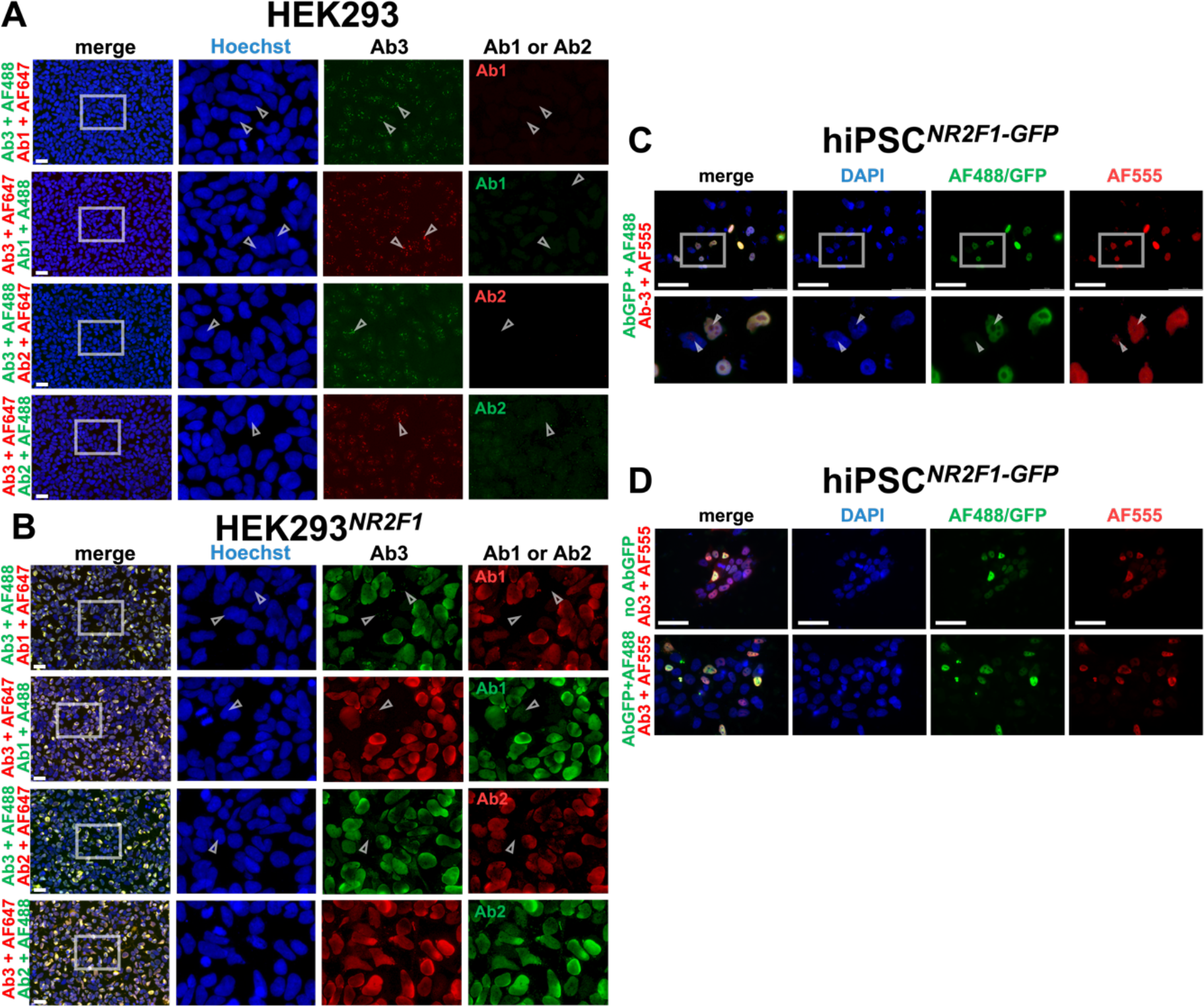
**(related to Figure 3): Double IF staining of overexpressed NR2F1 in different cell types by Ab3 with Ab1 or Ab2, and different 2°Abs**. 1°Abs and 2°Abs used are indicated on the left side of the pictures. A) Untransfected HEK293 cells co-stained with Ab3 and with either Ab1 or Ab2 in different combinations of adequate 2°Abs conjugated to AF555 (red) or AF647 (green). Arrowheads show nucleolar-like foci detected only in the Ab3 staining, regardless of which 2°Ab was used. B) In HEK293*^NR2F1^* Ab1, Ab2 and Ab3 detected nuclear NR2F1 localization. But Ab3 stained in addition the nucleolar-like foci (arrowheads), apparently only in cells not successfully transfected with NR2F1, which were negative for Ab1 and Ab2 staining. C-D) hiPSC*^NR2F1-GFP^*. C) Co- staining of Ab3 (red) with rabbit or mouse AbGFP and AF488 (green). Overview with the corresponding zoom-in below. Cells double-positive for GFP and Ab3 (yellow) display only a nuclear staining. GFP- negative (untransfected) cells show nucleolar-like foci stained by Ab3 (arrowheads). D) Cells stained with Ab1 only (upper line) and co-stained with mouse AbGFP and rabbit Ab1 (bottom line) show overlapping and nucleoplasmic staining only. Nucleoplasm (blue) was stained with (A-B) Hoechst or (C-D) DAPI. Scale bars: 50 µm.

**Supplementary Figure 6.**
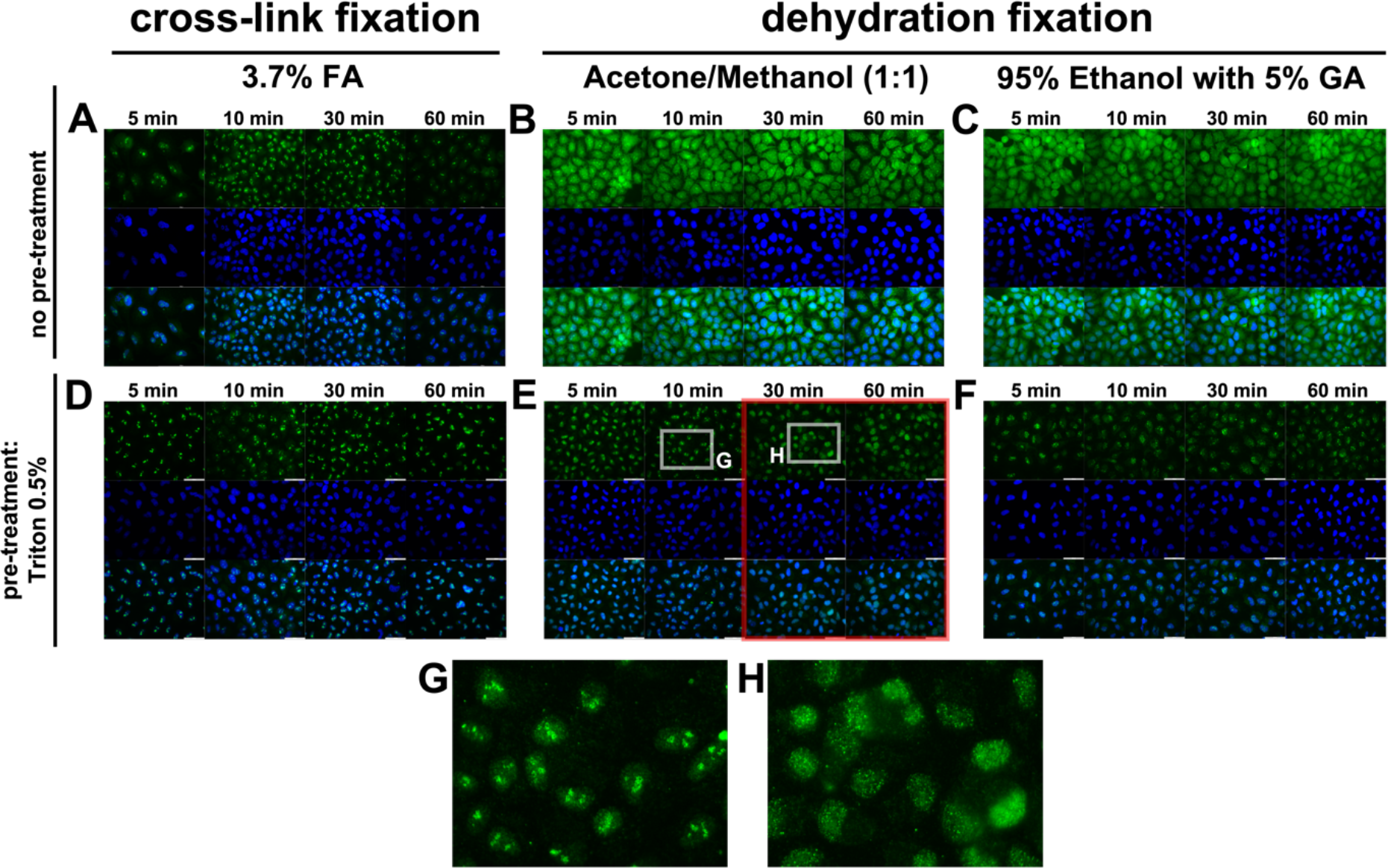
**(related to Figure 6): Testing different fixation times and methods for endogenous NR2F1 in HeLa cells by IF using Ab3.** Pre-treatment with 0.5% Triton for 0, (A-C) or 30 seconds (D-F) is indicated at the left side. Fixation times of 5, 10, 30 and 60 minutes for FA are in A, D, G), for Acetone/methanol mix in 1:1 ratio in B, E) and for 95% ethanol supplemented with 5% GA in C, F). White squares in E show zoom-in of G) nucleolar-like pattern of NR2F1 at 5-10 minutes of acetone/methanol fixation and H) and 30 minutes resulting in nucleoplasmic NR2F1 localization consistent with staining typically resulting from IF with all other Abs. Nucleoplasm (blue) was stained with DAPI. Scale bars: 50 µm.

**Supplementary Figure 7.**
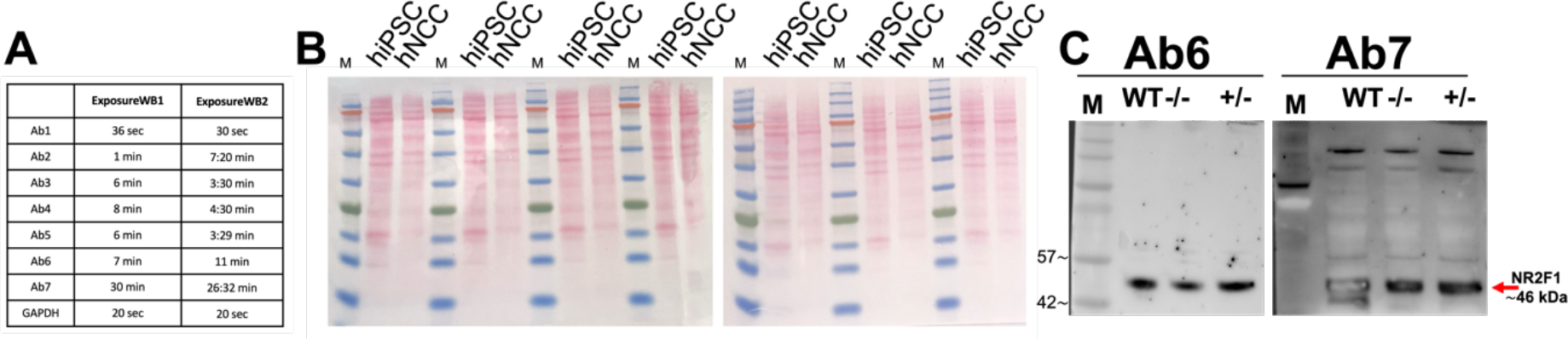
**(related to Figure 7E-F): Additional information about detection of NR2F1 in WB in *WT* hiPSC and hNCC as well as and hNCC*^-/-^* and hNCC*^+/-^*.** 20 µg per sample of RIPA lysates were loaded. A) Exposure times used for each antibody in both WBs showed in Figure 7B) Ponceau S staining from WB showed in Figure 7E, which demonstrates an efficient protein transfer onto the membranes. C) Membranes stained with Ab2 and Ab4 in WB2 that did not show a 46 kDa band in hNCC^-/-^ were stripped and stained again with Ab6 and Ab7, respectively. In hNCC^-/-^, both Ab6 and Ab7 stained a 46 kDa band corresponding to the molecular weight of NR2F1/NR2F2.

